# An interchange property for the rooted Phylogenetic Subnet Diversity on phylogenetic networks

**DOI:** 10.1101/2023.09.12.557317

**Authors:** Tomás M. Coronado, Gabriel Riera, Francesc Rosselló

## Abstract

Faith’s Phylogenetic Diversity (PD) on rooted phylogenetic trees satisfies the so-called strong exchange property that guarantees that, for every two sets of leaves of different cardinalities, a leaf can always be moved from the largest set to the smallest in such a way that the sum of the PD values does not decrease. This strong exchange property entails a simple polynomial-time greedy solution to the PD optimization problem on rooted phylogenetic trees. In this paper we obtain an exchange property for the rooted Phylogenetic Subnet Diversity (rPSD) on rooted phylogenetic networks of bounded level and reticulations’ in-degree, which involves a more complicated interchange of leaves. We derive from it a polynomial-time greedy solution to the rPSD optimization problem on rooted semibinary level-2 phylogenetic networks.

## 1 Introduction

Over the last few centuries, human activity has caused the destruction of natural habitats at an unprecedented pace, resulting in a major episode of biodiversity extinction [23]. Urgent action is required to combat extinction and preserve biodiversity, but there are challenges, including a lack of funding and uncertainties about conservation strategies. Consequently, there has been an increasing need to provide criteria for defining priorities and proposing variables that allow quantification of biodiversity.

The traditional approach to assessing biodiversity based on species counts, species richness, and number of endemic species has limitations. For instance, this type of data is so heterogeneous that it can be difficult to compare across different sites and times [13]. The approach based on lists of threatened species also has its drawbacks. For example, changes in the composition of these lists may represent changes in knowledge of species status rather than changes in the status itself [28]. Finally, measures of biodiversity based solely on species have been criticized for treating all species as equal, without regard to their functional roles in the ecosystem or their evolutionary history [10].

A feature of species that may influence their biodiversity value is their evolutionary distinctness. A species with few close living evolutionary relatives is considered more worthy of protection than a species with many close genetically and phenotypically similar relatives [24]. At the beginning of the 1990s, the qualitative value afforded to evolutionarily distinct species was replaced by quantitative measures of phylogenetic distinctness. One of the first measures of biodiversity based on phylogenetic information to appear in the literature was Faith’s *phylogenetic diversity*, PD [10]. The PD score of a set of species placed in the leaves of a phylogenetic tree is defined as the total weight (i.e., the sum of the branch lengths) of the spanning tree connecting the root and these leaves. In its original formulation, the branch lengths represented the number of changes in phenotypic characters, and PD measured the diversity of phenotypic characters in a set of species. In the current usual interpretation of phylogenetic trees, branch lengths represent evolutionary time, which is assumed to be positively correlated with character variation.

Since its introduction, PD has been widely studied and applied [27]. One of its most useful properties, both from the formal and the applicability point of view, is the possibility of efficiently finding and characterizing all subsets of species in a phylogenetic tree of a given size with maximal PD score by means of a very simple greedy algorithm [26, 33]; for instance, for a recent application to the analysis of SARS-CoV-2 phylogeny, see [40]. The basis of this result is the so-called *strong exchange property* stating that for every pair of sets of leaves *X, X*^*′*^ with |*X*| *>* |*X*^*′*^|, we can always move one leaf from *X* to *X*^*′*^ so that the sum of the PD scores does not decrease.

Faith’s PD is defined on evolutionary histories modelled by means of phylogenetic trees. But phylogenetic trees can only cope with speciation events due to mutations, where each species other than the universal common ancestor has only one parent in the evolutionary history (its parent in the tree). It is clearly understood now that other speciation events, which cannot be properly represented by means of single arcs in a tree, play an important role in evolution [9]. These are *reticulate events*, like genetic recombinations, hybridizations, or lateral gene transfers, where a species is the result of the interaction between two parent species. Reticulate processes are specially relevant in the description of microbial evolution [2, 31]. This has lead to the introduction of *phylogenetic networks* as models of phylogenetic histories that capture these reticulate events side by side with the classical mutations [19]. Faith’s PD has been recently extended both to split networks [32], a class of undirected graphs that generalize unrooted representing trees and do not describe evolutionary histories but simply evolutionary relationships, and to rooted phylogenetic networks [38]. In particular, in this last reference and in the later several generalizations of PD to rooted phylogenetic networks are proposed, being the most natural the *rooted Phylogenetic Subnet Diversity*, renamed *AllPaths-PD* in [4].

It has been proved that the PD optimization problem can be solved efficiently on *circular* split networks (a subclass of split networks widely used because they are the output of popular programs like PhyloNet [39] or Splitstree4 [18]) using integer programming [7, 32], as well as on the simplest class of non-tree rooted phylogenetic networks, the so-called *galled trees* [15], which model isolated hybridization events, by reducing it to set of linear size of minimum-cost flow problems [3, 4]. It has also been proved that the PD optimization problem is in general NP-hard on rooted phylogenetic networks [4] and on split networks [7], and it is easy to check that the strong exchange property does not hold even on galled trees [4].

In this paper we focus on the extension of the greedy optimization algorithm for PD on phylogenetic trees to a greedy algorithm for the optimization of rPSD on rooted phylogenetic networks. As we have mentioned, the greedy algorithm on phylogenetic trees is a consequence of the strong exchange property for PD that guarantees that the sum of the PD values of two sets of leaves of different cardinalities can always be increased by moving an element from the largest set to the smallest. That exchange property is no longer valid on phylogenetic networks. So, our first main contribution is a generalization to rPSD of that exchange property, through a more involved interchange of leaves than simply moving one leaf from one set to another.

Our exchange property then allows us to strengthen the result of Bordewich *et al* on galled trees, by proving that every rPSD-optimal set of *m* leaves in a galled tree is always obtained from an rPSD-optimal set of *m* − 1 leaves by either optimally adding a leaf or optimally replacing a leaf by a pair of leaves. Our exchange property also allows us to give a polynomial time greedy solution for the rPSD problem on semibinary level-2 networks and semi-3-ary level-1 networks, the next complexity level of rooted phylogenetic networks. On the negative side, we have not been able to deduce from it a greedy algorithm for semibinary level-3 or semi-4-ary level-1 networks and the problem for these more general classes remains open.

## 2 Preliminaries

### 2.1 Phylogenetic networks

Let Σ be a finite set of labels. By a *phylogenetic network* on Σ we understand a rooted directed acyclic graph where each node of in-degree ⩾ 2 has out-degree exactly 1 and whose *leaves* (i.e., its nodes of out-degree 0) are bijectively labeled in Σ [19]. A *phylogenetic tree* is simply a phylogenetic network without nodes of in-degree ⩾ 2. Let us point out here that, although the usual definition of phylogenetic tree and network forbids, for reconstructibility reasons, the existence of *elementary nodes*, that is, of nodes of in-degree and out-degree both equal to 1, we shall allow their existence in order to simplify some statements and proofs.

Let *N* be a phylogenetic network on Σ. We shall denote its *root* (i.e., its only node of in-degree 0) by *r* and its sets of nodes and arcs by *V* (*N*) and *E*(*N*), respectively, and we shall always identify its leaves with their corresponding labels. Given two nodes *u, v* in *N*, we say that *v* is a *child* of *u*, and also that *u* is a *parent* of *v*, when (*u, v*) ∈ *E*(*N*). A node in *N* is of *tree type*, or a *tree node*, when its in-degree is ⩽ 1, and a *reticulation* when its in-degree is ⩾ 2 (and hence, its out-degree is 1). An *exit reticulation* is a reticulation whose only child is a leaf: the rest of reticulations are called *internal*.

We shall say that *N* is *semi-d-ary* when all its reticulations have in-degree ⩽ *d*, and that *N* is *binary* when it is semi-binary and all its internal tree nodes have out-degree 2. *N* is said to be *tree-child* [6] when each internal node has a child of tree type: in particular, in a tree-child phylogenetic network the only child of every reticulation is a tree node.

We shall denote a (directed) path in *N* from a node *u* to a node *v* by *u* ⇝ *v*. The *intermediate* nodes of a path *u* ⇝ *v* are the nodes involved in it other than *u* and *v*. A *tree path* is a path with its origin and its intermediate nodes of tree type. A set of nodes *V*_0_ ⊆ *V* (*N*) is *independent* when it does not contain any pair of different nodes connected by a path.

For every *u, v* ∈ *V* (*N*), we say that *v* is a *descendant* of *u*, and also that *u* is an *ancestor* of *v*, when there exists a path *u* ⇝ *v*. In particular, every node is an ancestor, and a descendant, of itself. When *u* is an ancestor of *v* and *u* ≠ *v*, we shall say that *u* is a *strict ancestor* of *v* and that *v* is a *strict descendant* of *u*. We shall denote that *v* is a descendant of the end node of an arc *e* by *e* ≺ *v*.

The *cluster C*(*v*) ⊆ Σ of a node *v* ∈ *V* (*N*) is the set of (labels of) the descendant leaves of *v* and the *cluster C*(*e*) of an arc *e* ∈ *E*(*N*) is the cluster of its end node. For every *v* ∈ *V* (*N*), the *subnetwork of N rooted at v, N*_*v*_, is the subgraph of *N* induced by the set of all descendants of *v*. It is a phylogenetic network on *C*(*v*) with root *v*.

For every ∅= *X* ⊆ *V* (*N*), we shall denote by ↑_*N*_ *X*, or simply ↑*X* when there is no fear of amibiguity, the set of all nodes that are ancestors of nodes in ↑*X*. We shall often make the abuse of notation of using ↑*X* to denote also the subgraph of *N* induced by this set of nodes. In particular, given an edge *e* = (*u, v*) ∈ *E*(*N*), we shall write *e* ∈ ↑*X* to mean that *v* ∈ ↑*X*. As a graph, ↑*X* is a rooted directed acyclic graph with the same root *r* as *N*, and if, moreover, *X* ⊆ Σ, ↑*X* is a phylogenetic network on *X*.

A phylogenetic network *N* is *weighted* when it is endowed with a mapping *w* : *E*(*N*) → ℝ_+_ that assigns a weight *w*(*e*) ⩾ 0 to every arc *e*. The *total weight* of a subgraph of a weighted phylogenetic network is the sum of the weights of all arcs in the subgraph. In particular, the weight *w*(*u* ⇝ *v*) of a path *u* ⇝ *v* is the sum of the weights of its arcs.

A subgraph of an undirected graph is *biconnected* when it is connected and it remains connected after removing any node from it and all arcs incident to this node. A subgraph of a phylogenetic network *N* is *biconnected* when it is so in the undirected graph underlying *N*. Every node and every arc in *N* are biconnected subgraphs, and every biconnected subgraph of *N* with more than 2 nodes contains at least one reticulation.

A *biconnected component* of *N* is a maximal biconnected subgraph. Every biconnected component *C* of a phylogenetic network has one, and only one, node that is ancestor of all nodes in *C*. When *C* has more than 2 nodes, we shall call this node the *split node* of *C*. Moreover, if *C* has more than 2 nodes, every node in *C* with no child inside *C* is a reticulation (should it be of tree type, removing its parent would disconnect *C*). We shall call such nodes the *exit reticulations* of *C*, and the rest of the reticulations of *C, internal*. We shall also use the term *blob* to mean a biconnected component with more than 2 nodes of a phylogenetic network. As we have just seen, every blob contains some reticulation.

A phylogenetic network is *level-k* [21] when every biconnected component contains at most *k* reticulations. Thus, a level-0 network is a phylogenetic tree. A semibinary level-1 network is also called a *galled tree* [15]. Each one of the blobs of a galled tree, called in this case *galls*, consists of two paths going from the same tree node to the same reticulation without any intermediate node in common. The phylogenetic network in Fig. 1 is a galled tree. To simplify the language, we shall call a *semi-d-ary k-blob* any biconnected component with *k* ⩾ 1 reticulations of a semi-*d*-ary phylogenetic network.

**Figure 1:**
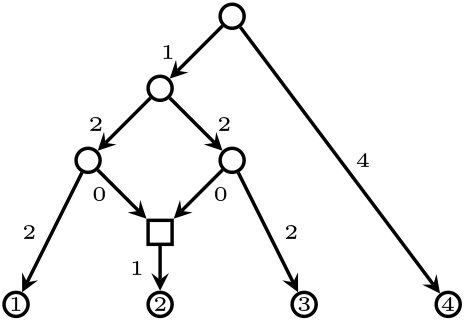
A weighted phylogenetic network. The tree nodes are represented by circles, the reticulation by a square, and the arcs’ labels represent their weights.

We end this subsection with a lemma for which we have not found any suitable reference in the literature even in the semibinary (*d* = 2) case.

#### Lemma 1.

*Let N be a blob with set of internal reticulations* 𝒮 *and set of exit reticulations* ℰ. *Let* 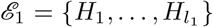 *and let* 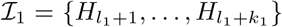 *be the set of internal reticulations in* ↑_*N*_ ℰ_1_ *\* ↑_*N*_ (ℰ *\* ℰ_1_). *For each i* = 1, …, *l*_1_ + *k*_1_, *let d*_*i*_ = deg_*in*_ *H*_*i*_. *Then, for every independent set of nodes V* ⊆ ↑_*N*_ ℰ_1_ *\* ↑_*N*_ (ℰ *\* ℰ_1_),

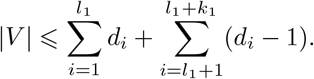

*Proof*. We shall prove the thesis by double induction on *l*_1_ = |ℰ_1_| and *k* = |𝒮|. The case when *l*_1_ = 0 is obvious, because then ↑_*N*_ ℰ_1_ *\* ↑_*N*_ (ℰ *\* ℰ_1_) = ∅.

To prove the general inductive step, from *l*_1_ − 1 to *l*_1_, we begin with the case when *k* = 0. So, let *N* be a blob without internal reticulations, let *E* be its set of exit reticulations, and let 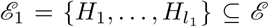 with *l*_1_ ⩾ 1. Let us assume, as induction hypothesis, that the thesis in the statement is true for all blobs *N* ^*′*^ without internal reticulations and subsets of exit reticulations 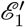 of cardinality 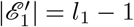.

Take a node 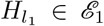 and let 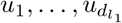 be its parents. For each 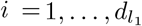, let *v*_*i*_ be the lowest ancestor of *u*_*i*_ that has some descendant (exit) reticulation other than 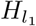, and consider the path *v*_*i*_ ⇝ *u*_*i*_. Concatenating each such path with the corresponding arc 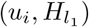, we obtain 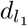 different paths 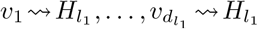 ending in 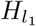 : observe that the nodes 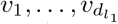 need not be different, but each such path ends in a different arc 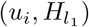.

Let *N* ^*′*^ be the directed graph obtained by removing from *N* the reticulation 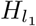 and the intermediate nodes of the paths 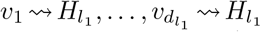, together with the arcs incident to them (if some path is simply an arc 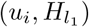, we remove the arc). The set of exit reticulations of *N* ^*′*^ is 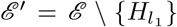; take 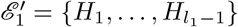. Since *N* did not have internal reticulations, neither does *N* ^*′*^. Then, by the induction hypothesis, for every independent set of nodes *V* in 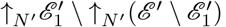,

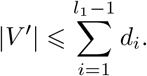

Now, any such independent set of nodes *V* ^*′*^ in 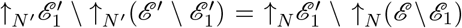 can be enlarged to an independent set of nodes *V* in ↑_*N*_ ℰ_1_ \↑_*N*_ (ℰ *\*ℰ_1_) by adding at most 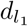 nodes, one inside each path 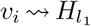. Conversely, if we remove from an independent set of nodes *V* in *N* its nodes belonging to the paths 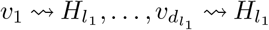 (and by the independence condition each such path will contain at most one element of *V* and therefore we remove in this way at most 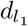 nodes), we obtain an independent set of nodes *V* ^*′*^ in *N* ^*′*^. Then, since 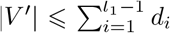, any independent set of nodes in ↑_*N*_ ℰ_1_ \↑_*N*_ (ℰ *\*ℰ_1_) has cardinality at most 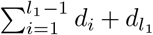. This proves this inductive step when *k* = 0.

Let us prove now, for any fixed *l*_1_ *>* 0, the inductive step from *k* − 1 to *k*. So, assume that the thesis in the statement is true for all blobs *N* ^*′*^ with *k* − 1 internal reticulations and subsets 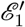 of exit reticulations of cardinality 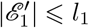, and let *N* be a blob with *k* internal reticulations, let ℰ be its set of exit reticulations, and let 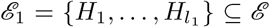 with *l*_1_ ⩾ 1. Let 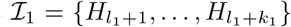 be the set of internal reticulations in ↑_*N*_ ℰ_1_ \ ↑_*N*_ (ℰ *\*ℰ_1_).

Let *H* be an internal reticulation with no reticulate ancestor and let *u*_1_, …, *u*_*d*_ be its parents. For each *i* = 1, …, *d*, let *v*_*i*_ be the lowest ancestor of *u*_*i*_ with some reticulation descendant that is not a descendant of *H*. Concatenating each path *v*_*i*_ ⇝ *u*_*i*_ with the corresponding arc (*u*_*i*_, *H*), we obtain *d* different paths *v*_1_ ⇝ *H*, …, *v*_*d*_ ⇝ *H* ending in *H*: as before, observe that the nodes *v*_1_, …, *v*_*d*_ need not be different, but each such path ends in a different arc (*u*_*i*_, *H*). Let *N* ^*′*^be the directed graph obtained by removing the intermediate nodes in all paths *v*_*i*_ ⇝ *H* except for one path *v*_1_ ⇝ *H*, together with the arcs incident to them; if *v*_*i*_ = *u*_*i*_, we simply remove the arc (*u*_*i*_, *H*). The blob *N* ^*′*^ still has the same set of exit reticulations ℰ as *N*, but it has *k* − 1 internal reticulations because *H* has become an elementary tree node in *N* ^*′*^. Moreover, if we denote by 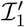 the set of internal reticulations in ↑_*N′*_ ℰ _1_ \ ↑_*N*′_ (ℰ *\*ℰ _1_), then 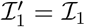 if *H* ∉ 𝒮_1_ and 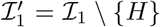 if *H* ∈ 𝒮_1_. If this last case happens, let us assume without any loss of generality that 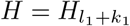 and hence that 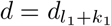.

Therefore, by the induction hypothesis, for every independent set of nodes *V* ^*′*^ in ↑_*N*′_ ℰ_1_ \ ↑_*N*′_ (ℰ *\* ℰ_1_)

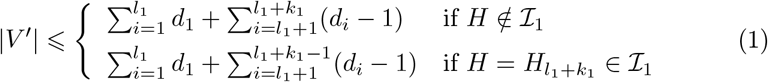

Now, notice that any independent set of nodes *V* ^*′*^ in *N* ^*′*^ can be enlarged to an independent set of nodes *V* in *N* by adding at most one node inside each one of the *d* − 1 removed paths *v*_*i*_ ⇝ *H*. Conversely, if we remove from an independent set of nodes *V* in *N* its nodes that are intermediate in the *d* − 1 removed paths *v*_*i*_ ⇝ *H* (and by the independence condition each such path will contain at most one element of *V* and therefore we are removing in this way at most *d* nodes from *V*), we obtain an independent set of nodes *V* ^*′*^ in *N* ^*′*^. Then:

- If *H* ∉ ↑_*N*_ ℰ_1_ \ ↑_*N*_ (ℰ *\* ℰ_1_), any maximal independent set of nodes ↑_*N*′_ ℰ_1_ \ ↑_*N*′_ (ℰ *\*ℰ_1_) is also a maximal independent set of nodes in ↑_*N*_ ℰ_1_ \↑_*N*_ (ℰ *\*ℰ_1_). Then, by Eqn. (1), 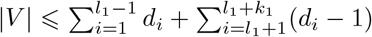 for every maximal independent set of nodes *V* in ↑_*N*_ ℰ_1_ \ ↑_*N*_ (ℰ *\* ℰ_1_).
- If 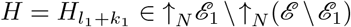 any maximal independent set of nodes in ↑_*N*′_ ℰ_1_ \ ↑_*N*′_ (ℰ *\*ℰ_1_) can be enlarged to a maximal independent set of nodes in ↑_*N*_ ℰ_1_ \ ↑_*N*_ (ℰ *\*ℰ_1_) by adding at most 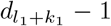 nodes. Then, by Eqn. (1), 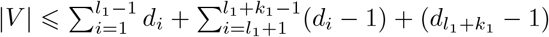 for every maximal independent set of nodes *V* in ↑_*N*_ ℰ_1_ \ ↑_*N*_ (ℰ *\*ℰ_1_).

This finishes the proof of the inductive step.

#### Corollary 2.

*Let N be a semi-d-ary blob*, ℰ *its set of exit reticulations*, 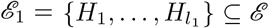, *and k*_1_ *the number of internal reticulations of N belonging to* ↑_*N*_ ℰ_1_ \ ↑_*N*_ (ℰ *\*ℰ_1_). *Then, for every independent set of nodes V in* ↑_*N*_ ℰ_1_ \ ↑_*N*_ (ℰ *\*ℰ_1_),

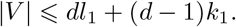

*In particular, if N has k internal reticulations*, |*V* | ⩽ *dl*_1_ + (*d* − 1)*k*.

The constructions explained in the proof of Lemma 1 easily show that, for every *d, k, l, l*_1_ with *d* ⩾ 2 and *k* ⩾ *l* ⩾ *l*_1_ ⩾ 1, there are semi-*d*-ary *k*-blobs with *l* exit reticulations and subsets *E*_1_ of *l*_1_ exit reticulations containing an independent set of nodes *V* in ↑_*N*_ *E*_1_ \↑_*N*_ (*E \E*_1_) with *l*_1_*d* + (*k* − *l*)(*d* − 1) elements: cf. Fig. 2.

**Figure 2:**
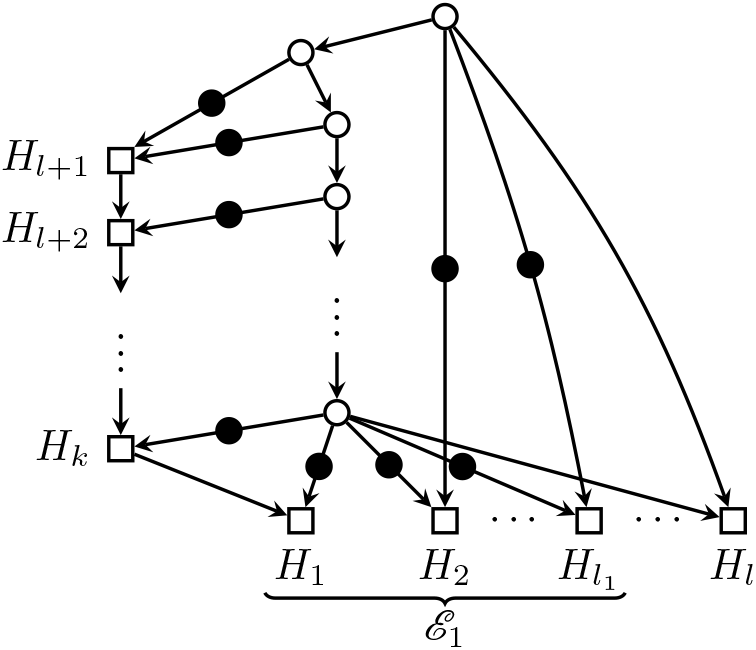
A semibinary *k*-blob *G* with *l* exit reticulations, a distinguished subset ℰ_1_ of *l*_1_ exit reticulations, and an independent set of nodes (represented by filled circles) in ↑_*G*_ℰ_1_ ∩ (↑_*G*_(ℰ \ ℰ1))^*c*^ reaching the upper bound in Corollary 2 for *d* = 2.

#### Remark 3.

By Lemma 1, and with the notations in its statement, if *N* is a blob without internal reticulations and *V* is an independent set of nodes of ↑_*N*_ *E*_1_ \ ↑_*N*_ (*E \ E*_1_), then 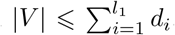. A close analysis of the proof of that lemma easily shows that the upper bound 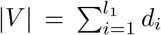 is achieved at the sets *V* containing, for every *H*_*i*_ ∈ *ℰ*_1_, exactly *d*_*i*_ nodes whose only reticulate descendant is *H*_*i*_. Of course, such sets do not always exist: for instance, when, *N* contains a node that is a parent of two different exit reticulations.

### 2.2 The rooted Phylogenetic Diversity on phylogenetic trees

Given a finite set Σ, we shall denote henceforth its set of subsets by 𝒫 (Σ) and, for every *k* ⩾ 0, the set of all its subsets of cardinality *k* by 𝒫_*k*_(Σ).

Given a weighted phylogenetic tree *T* on Σ, Faith’s *rooted Phylogenetic Diversity* [10] is the set function rPD_*T*_ : 𝒫 (Σ) → ℝ_+_ sending each *X* ⊆ Σ to the *total weight* of ↑_*T*_*X*:

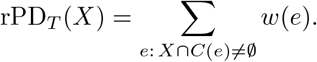

This rooted Phylogenetic Diversity rPD_*T*_ on phylogenetic trees satisfies the following *strong exchange property*, introduced in [33] for unrooted phylogenetic trees: for every phylogenetic tree *T* on Σ and for every *X, X*^*′*^ ⊆ Σ such that |*X*^*′*^| *<* |*X*|, there exists some *a* ∈ *X \ X*^*′*^ such that

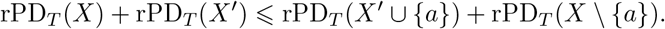

For a proof of this fact in the rooted case, see [34, §6.4.1].

This strong exchange property for rPD_*T*_ is the key ingredient in the proof that the following simple greedy algorithm produces, for every *k* ⩾ 1, the family ℳ_*k*_ of all rPD_*T*_ *-optimal* subsets of Σ of cardinality *k*, that is, of all sets of *k* leaves with maximum rPD_*T*_ :

#### Algorithm 1: Greedy for phylogenetic trees

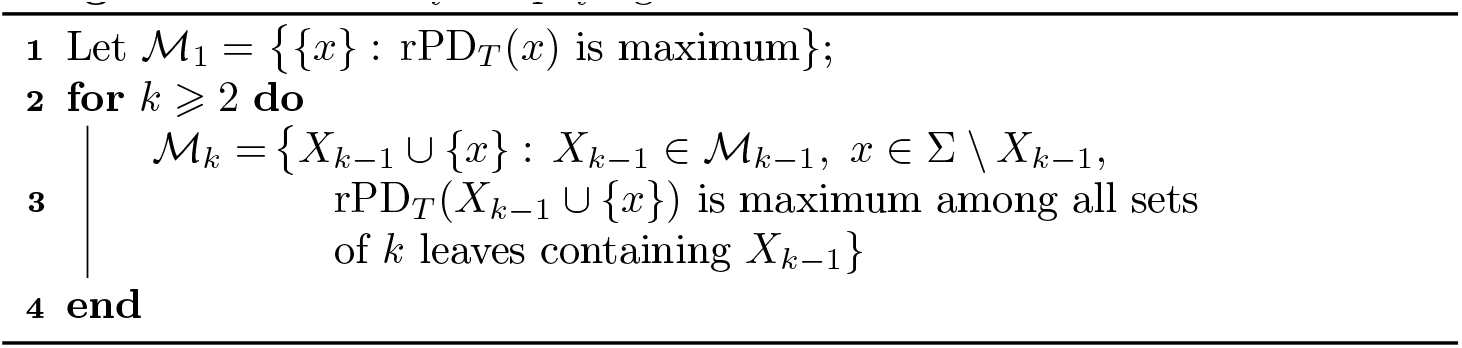

For a proof in the unrooted case, see [33]; the proof in the rooted case is similar (cf. [34, §6.4.1]).

In particular, given a weighted phylogenetic tree *T* on Σ, this provides a polynomial solution to the problem of finding the maximum rooted Phylogenetic Diversity value among all members of 𝒫_*k*_(Σ), and a member of 𝒫_*k*_(Σ) reaching this maximum.

## 3 The rooted Phylogenetic Subnet Diversity

K. Wicke and M. Fischer [38] proposed several generalizations of Faith’s rooted Phylogenetic Diversity function to weighted phylogenetic networks. One of them, and possibly the most straightforward, is the *rooted Phylogenetic Subnet Diversity* : the set function rPSD_*N*_ : 𝒫 (Σ) → ℝ_+_ sending each *X* ⊆ Σ to the total weight of ↑*X*:

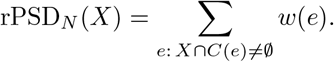

It is clear that if *N* is a phylogenetic tree, then rPSD_*N*_ = rPD_*N*_.

### Example 4.

On the phylogenetic network *N* depicted in Figure 1,

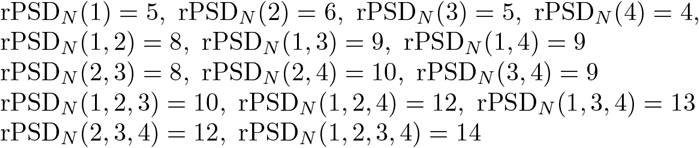

For every weighted phylogenetic network *N* on Σ, its set function rPSD_*N*_ is:

i. *Monotone increasing*: For every *X* ⊆ *Y* ⊆ Σ,

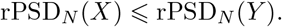
ii. *Submodular* : For every *X* ⊆ *Y* ⊆ Σ and for every *a* ∈ Σ *\ Y*,

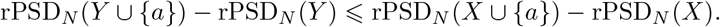
iii. *Subadditive*: For every *X, Y* ⊆ Σ,

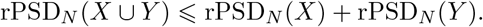

(i) and (iii) are clear. As to (ii), it is proved in [4, Lemma 4.3].

On the negative side, rPSD_*N*_ needs not satisfy the strong exchange property, even for the simplest non-tree networks *N*. Indeed, consider again the weighted galled tree *N* with a single reticulation depicted in Figure 1. Take *X* = *{*1, 3, 4*}* and *X*^*′*^ = *{*2, 4*}*. Then

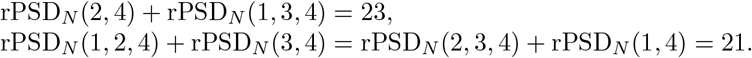

Therefore, there is no *a* ∈ *X \ X*^*′*^ such that

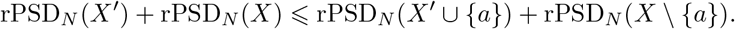

As a consequence, a rPSD_*N*_ -optimal set of cardinality *k* need not contain any rPSD_*N*_ -optimal set of cardinality *k* − 1. Consider again the galled tree depicted in Figure 1. Its set of 2 labels with largest rPSD_*N*_ value is *{*2, 4*}*, while its set of 3 labels with largest rPSD_*N*_ value is *{*1, 3, 4*}*, which is not obtained by expanding *{*2, 4*}*.

So, Algorithm 1 cannot be used to produce rPSD-optimal sets of a given cardinality as it stands. Actually, Bordewich *et al* prove in [4, Thm. 4.1] that, given a phylogenetic network *N* on Σ and an integer *k*, the problem of finding the maximum rPSD_*N*_ value on 𝒫 _*k*_(Σ) is NP-hard. On the positive side, these authors prove in [4, Prop. 4.5] that this problem can be solved in polynomial time on binary galled trees.

## 4 A general exchange property

Let Σ be a finite set and *W* : 𝒫 (Σ) → ℝ _+_ a function. Given *X, X*^*′*^ ⊆ Σ such that |*X*^*′*^| *<* |*X*|, a *W -improving pair* for *X, X*^*′*^ is a pair of sets (*A, B*) with *A* ⊆ *X \ X*^*′*^ and *B* ⊆ *X*^*′*^ *\ X* such that |*B*| *<* |*A*| and

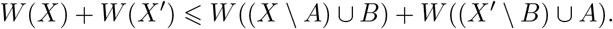

Let 𝒮 ⊆ (*A, B*) ∈ 𝒫 (Σ)^2^ : |*B*| *<* |*A*|. We shall say that *W* : 𝒫 (Σ) → ℝ _+_ satisfies the *exchange property with respect to* 𝒮 when every pair of sets *X, X*^*′*^ ⊆ Σ with |*X*^*′*^| *<* |*X*| has a *W* -improving pair in 𝒮.

With these notations, Steel’s *strong exchange property* for phylogenetic trees mentioned in §2.2 says that, for every weighted phylogenetic tree *T* on Σ, rPD_*T*_ : 𝒫 (Σ) → ℝ_+_ satisfies the exchange property with respect to

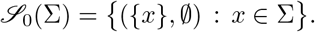

As we have seen, galled trees need not satisfy Steel’s strong exchange property. The main result in this section, Theorem 6, says that every semi-*d*-ary level-*k* phylogenetic network on Σ satisfies the exchange property with respect to the following larger family of pairs of subsets 𝒮_*k,d*_(Σ) whose description only depends on *k* and *d*: when *k* = 1,

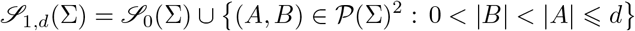

and, if *k* ⩾ 2,

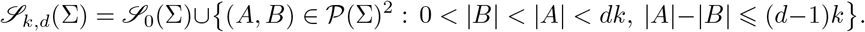

Notice that the only pairs (*A, B*) ∈ 𝒮_*k,d*_(Σ) with *B* = ∅ have |*A*| = 1. Given *k* and *d*, the cardinalities of these families of sets are polynomial in |Σ| = *n*: |𝒮_0_(Σ)| = *n* and

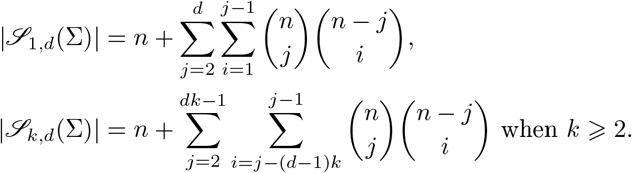

To simplify the notations, given *X* ⊆ Σ, *S* ⊆ *X* and *T* ⊆ Σ \*X*, we shall denote (*X* \ *S*) ∪ *T* by *τ*_*S,T*_ (*X*).

The next lemma extracts a key technical step in the proof of Theorem 6. More precisely, we prove an analog of the exchange property for the nodes of a blob, labelled as *X* or *X*^*′*^ with multiplicities which represent the number of descendant leaves belonging to *X* or *X*^*′*^ in Theorem 6. Although that theorem deals with sets of leaves and edge weights, this lemma is completely local to the blob and independent of any weighting.

We use in it some standard notations for multisets *X*: *m*_*X*_ (*v*) denotes the multiplicity of an element *v* in *X*, Supp *X* denotes the *support* of *X*, that is, the set of elements *v* such that *m*_*X*_ (*v*) *>* 0, and the cardinality of *X* is |*X*| = Σ_*x*∈Supp *X*_ *m*_*X*_ (*v*). We shall also use the notation *τ*_*S,T*_ (*X*) = (*X* \ *S*) ∪ *T* when *X, S, T* are multisets with *S* ⊆ *X* and Supp *T* ⊆ Σ \ Supp *X*.

This lemma also uses some basic properties of ↑-notation. Some simple results to facts into consideration in this regard are that, for any *A, B*, ↑ (*A* ∪ *B*) = ↑*A* ∪ ↑*B* and ↑*A \* ↑*B* ⊆↑ (*A\ B*), and that if *A* ⊆ *B*, then ↑*A* ⊆ ↑*B*. Moreover, given a multiset *A*, we define ↑*A* as ↑Supp *A*, without taking into account the multiplicities of the elements of *A*.

### Lemma 5.

*Let B be a semi-d-ary level-k blob and X, X*^*′*^ *two multisets of nodes of B satisfying the following properties:*

i. |*X*^*′*^| *<* |*X*|.
ii. *For each v* ∈ *V* (ℬ), *if m*_*X*′_ (*v*) *< m*_*X*_ (*v*), *then m*_*X*_ (*v*) = 1.
iii. *Each exit reticulation of* ℬ *belongs to X or X*^*′*^.

*Then there exist a set A and a multiset B of nodes of* ℬ *such that:*

1. *B* = ∅ *and* |*A*| = 1 *or*

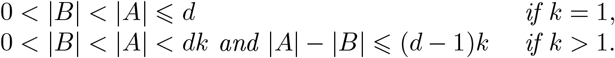
2. *A* ⊆ Supp *X \* Supp *X*^*′*^.
3. Supp *B* ⊆ Supp *X*^*′*^ \ Supp *X and m*_*B*_ *is the restriction of m*_*X*′_ *to* Supp *B*.
4. ↑*X* ∩ ↑*X*^*′*^ ⊆ ↑*τ*_*A,B*_(*X*) ∩ ↑*τ*_*B,A*_(*X*^*′*^).

*Proof*. First, we introduce some notation.

- 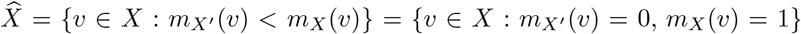. Observe that 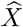 is a set: *X* 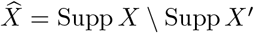.
- 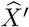 is the multiset with support *{v* ∈ *X*^*′*^ : 0 = *m*_*X*_ (*v*) *< m*_*X*′_ (*v*)*}* = Supp *X*^*′*^ *\* Supp *X* and multiplicities from *X*^*′*^.
- *ℰ* is the set of exit reticulations of ℬ. In addition, by (iii), we classify *E* into three disjoint subsets: 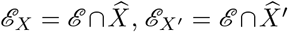 and *ℰ* _*X,X*′_ = *ℰ* ∩ *X* ∩ *X*^*′*^.
- For each exit reticulation *H* ∈ *ℰ*, let *D*_*H*_ = ↑*H \* ↑ (*ℰ \* {*H*}). This is, *D*_*H*_ consists of the ancestors of *H* that are not ancestors of any other exit reticulation. Observe that the sets *D*_*H*_, with *H* ∈ *ℰ*, are pairwise disjoint.

Using these notations, we have the following sequence of inequalities:

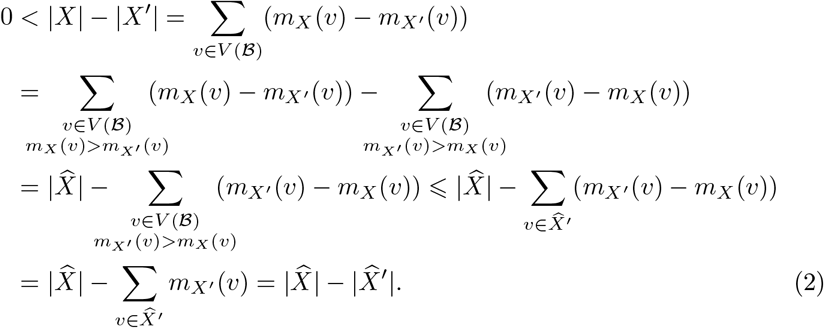

Now there are three cases to consider, in all of them we shall choose a subset 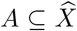 and a submultiset 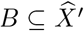 with *m*_*B*_ the restriction on 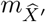 to Supp *B*, and hence (2) and (3) will hold true.

a. Assume that there exists some 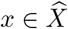 with a strict descendant in *X*. In this case, take *A* = *{x}* and *B* = ∅. Then (1) is clearly satisfied and

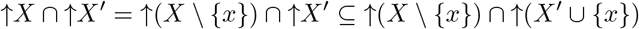

because ↑*X* = ↑(*X* \ *{x}*) since *x* ∈ ↑(*X \ {x}*).
b. Assume that no 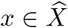 has any strict descendant in *X* and that ℰ _*X*′_ = ∅. This implies that ℰ = ℰ _*X*_ ∪ ℰ _*X,X*′_ ⊆ *X* and that 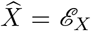, as any 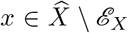 would have some descendant in ℰ ⊆ *X*. Then, Eqn. (2) implies

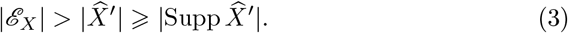

Using this fact, we can now prove the existence of some *H*_0_ ∈ ℰ _*X*_ satisfying ↑*H*_0_ ∩ ↑*X*^*′*^ ⊆ ↑(*X \ {H*_0_*}*):
  - First, we prove the existence of some *H*_0_ ∈ ℰ _*X*_ such that 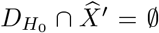 by contradiction. Indeed, if 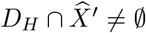 for every *H* ∈ ℰ _*X*_, then we can choose for each *H* ∈ ℰ _*X*_ some element in 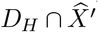. Since the sets *D*_*H*_ are pairwise disjoint, this provides ℰ _*X*_ different elements in Supp 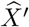, which contradicts Eqn. (3). which does not exist.
  - So, let *H*_0_ ∈ ℰ _*X*_ be such that 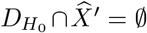. Then, 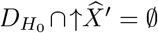 because any node 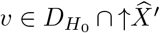 would have all its descendants contained in 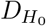 and then any descendant of *v* in 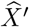 would be an element of 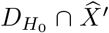, which does not exist.
  - Now let *v* ∈ ↑*H*_0_ ∩ ↑*X*^*′*^. Then, either *v* ∈ 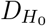or *v* ∈ ↑(ℰ \ *{H*_0_*}*). In the first case, from the previous bullet point we have 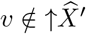, so

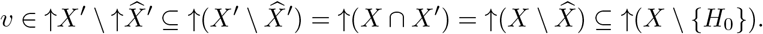

In the second case we already have *v* ∈ ↑(ℰ \ *{H*_0_*}*) ⊆ ↑(*X* \ *{H*_0_*}*). Take *A* = *{H*_0_*}* and *B* = ∅ so that (1) is satisfied. Then

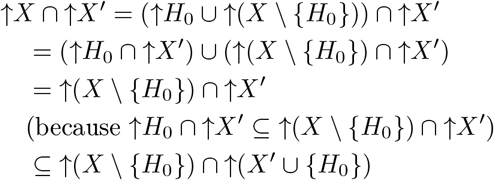

which proves (4).
c. Assume that no 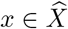 has any strict descendant in *X* and that ℰ _*X*′_≠∅. Then, 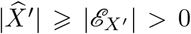 and, by Eqn. (2), 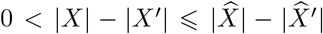. Let *l*_0_ = |ℰ _*X*_ |, *l*_1_ = |ℰ _*X*′_ | and *l*_2_ = |ℰ _*X,X*′_ |. Then, by (iii), *l*_0_ + *l*_1_ + *l*_2_ = |ℰ | ⩽ *k* and, by the first assumption, 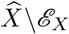 is an independent set of nodes of ↑ℰ _*X*′_*\* ↑ (ℰ *\* ℰ _*X*′_). Thus by Lemma 1, and since *d* ⩾ 2, we have

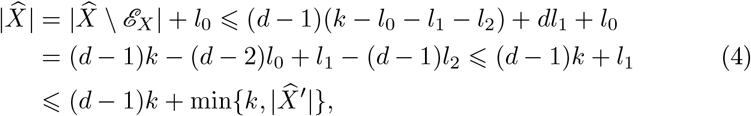

using that *l*_1_ ⩽ |ℰ | ⩽ *k* and 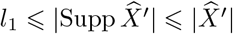. In particular,

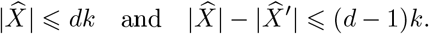

Let us now further distinguish two cases.
  (c.1) First, assume that 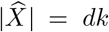. In this case, all inequalities in the sequence (4) as well as the inequalities *l*_1_ ⩽ℰ ⩽ *k* are equalities. This implies that ℰ = ℰ _*X*′_ and the blob ℬ has no internal reticulation. Moreover, since the first inequality in (4) is an equality, 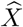 reaches the maximum number of possible independent nodes in ↑ℰ _*X*′_. Then, as noted in Remark 3, it must happen for each *H* ∈ *E* that deg_*in*_(*H*) = *d* and 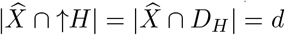. Now, since 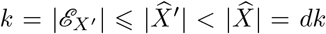, there must exist some *H*_0_ ∈ *E*_*X*′_ with *m*_*X*′_ (*H*_0_) *< d*. Take then 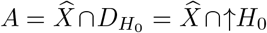 and the multiset *B* with Supp *B* = {*H*_0_} and *m*_*B*_(*H*_0_) = *m*_*X*′_ (*H*_0_). We have that 0 *<* |*B*| *<* |*A*| = *d* and hence the pair (*A, B*) satisfies condition (1) in the statement. As to condition (4),

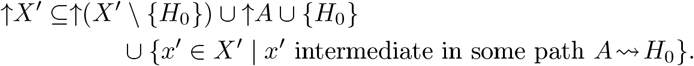

Now, *H*_0_ *∉* ↑*X* and, by assumption, the elements of *A* have no strict descendant in *X*, which implies

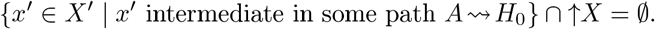

Moreover, since *A* ⊆ ↑*H*_0_, we have that ↑*X* ⊆ ↑((*X* \ *A*) ∪ *{H*_0_*}*). Therefore

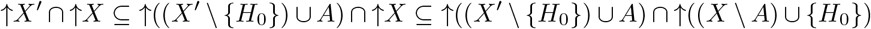

which proves (4).
  (c.2) Assume now that 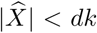. Take 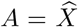 and 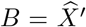. Then (*A, B*) satisfy the required cardinality conditions in (1) and

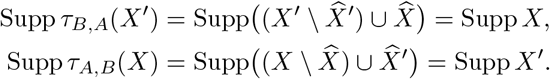

and hence ↑*X*^*′*^ ∩ ↑*X* = ↑*τ*_*A,B*_(*X*) ∩ ↑*τ*_*B,A*_(*X*^*′*^), which implies (4).

### Theorem 6.

*If N is a semi-d-ary level-k weighted phylogenetic network on* Σ, rPSD_*N*_ *satisfies the exchange property with respect to* 𝒮_*k,d*_(Σ).

*Proof*. As we have already mentioned, the case *k* = 0 is Steel’s strong exchange property for phylogenetic trees [34, §6.4.1]. So, we shall focus on the case *k* ⩾ 1.

Without any loss of generality, we can assume that every tree node in *N* is at most bifurcating (recall that we do not forbid out-degree 1 tree nodes in our networks). Indeed, let first *N* ^*′*^ be the phylogenetic network obtained from *N* as follows: for every node *v* that is the split node of more than one blob and for each such blob rooted at *v*, add a new split node *v*_*i*_ to the blob and a new edge (*v, v*_*i*_) with weight 0. *N* ^*′*^ is still semi-*d*-ary and level-*k*, no node in it is the split node of more than one blob, and rPSD_*N*_ (*Z*) = rPSD_*N*′_ (*Z*) for every *Z* ⊆ Σ. Now, let *N* ^*′′*^ be the phylogenetic network obtained from *N* ^*′*^ as follows: for every tree node *v* with *k* ⩾ 3 children *v*_1_, …, *v*_*k*_, replace in *N* the subgraph supported non {*v, v*_1_, …, *v*_*k*_ } by a bifurcating tree with root *v* and leaves *v*_1_, …, *v*_*k*_ and all its edges except those ending in *v*_1_, …, *v*_*k*_ of weight 0: the edge ending in each *v*_*i*_ inherits the original weight of (*v, v*_*i*_); if any node *v*_*i*_ had any entering edges other than (*v, v*_*i*_), we keep them with their weights. Since *v* was the split node of at most one blob, no blob increases its level from *N* ^*′*^ to *N* ^*′′*^, and therefore *N* ^*′′*^ is still semi-*d*-ary and level-*k*, and rPSD_*N*′′_ (*Z*) = rPSD_*N*′_ (*Z*) = rPSD_*N*_ (*Z*) for every *Z* ⊆ Σ.

So, in the rest of this proof we shall suppose that *N is at-most-bifurcating*, in the sense that the out-degree of each tree node is at most 2, and in particular that no node in *N* is the split node of more than one blob. To simplify the notations, for every node *v* in *N* we shall denote its cluster *C*_*N*_ (*v*) by *C*(*v*).

We shall proceed by induction on the number *α* of edges of the network. A phylogenetic network with *α* = 0 is a phylogenetic tree consisting of a leaf, where the stated exchange property trivially holds. Now, let *N* be an at-most-bifurcating semi-*d*-ary level-*k* phylogenetic network on Σ with *α* ⩾ 1 edges, and let us suppose that the thesis in the statement is true for all at-most-bifurcating semi-*d*-ary level-*k* phylogenetic networks with less than *α* edges.

Let *X, X*^*′*^⊆ Σ with |*X*^*′*^| *<* |*X*|. If |*X*| = 1 the exchange property is trivially satisfied taking *A* = *X* and *B* = *X*^*′*^ = ∅, so we assume from now on that |*X*| ⩾ 2. Now consider the tree of blobs *T* of *N* [16], obtained by collapsing each blob in *N* into its split node. Then, *T* is a phylogenetic tree on Σ with the same root *r* as *N* and, for every node *v* in *T, C*_*T*_ (*v*) = *C*(*v*). Since |*X*^*′*^| *<* |*X*| and |*X*| ⩾ 2, the set of nodes *v* in *T* such that |*X*^*′*^ ∩ *C*(*v*)| *<* |*X* ∩ *C*(*v*)| and 1 *<* |*X* ∩ *C*(*v*)| is not empty: it contains at least the root *r*.

There are four cases to consider.

a. Assume that *T* contains some node *v*_0_ different from the root *r* such that |*X* ∩ *C*(*v*_0_) | *>* |*X*^*′*^∩ *C*(*v*_0_) | and |*X* ∩ *C*(*v*_0_) | *>* 1. In particular, *v*_0_ is a tree node of *N* such that the edge *e*_0_ = (*v*_1_, *v*_0_) ending in it is a cut edge. Let 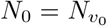 and let *N*_1_ be the network obtained from *N* by removing 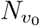 and the edge *e*_0_ and, if *v*_1_ is a reticulation node, appending to it a dummy leaf child (not labelled in Σ) through an edge of weight 0. Both *N*_0_ and *N*_1_ are at-most-bifurcating, semi-*d*-ary, level-*k*, and have less than *α* edges, and therefore by assumption they satisfy the thesis in the statement. Now, for every *Z* ⊆ Σ,

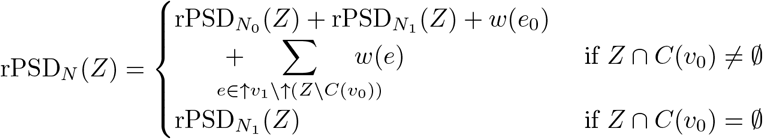

(Throughout this proof, given a network *N* ^*′*^ with set of leaves Σ^*′*^ and a set *Z*, we write rPSD_*N*′_ (*Z*) to denote actually rPSD_*N*′_ (*Z* ∩ Σ^*′*^). So, for instance, 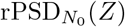 and 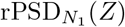 in the expression above actually mean 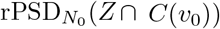 and 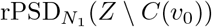, respectively.) Now, since |*X* ∩ *C*(*v*_0_)| *>* |*X*^*′*^ ∩ *C*(*v*_0_)|, by the induction hypothesis there exist *A* ⊆ (*X \ X*^*′*^) ∩ *C*(*v*_0_) and *B* ⊆ (*X*^*′*^ *\ X*) ∩ *C*(*v*_0_) such that (*A, B*) ∈ *𝒮*_*k,d*_(*C*(*v*_0_)) ⊆ *𝒮*_*k,d*_(Σ) and

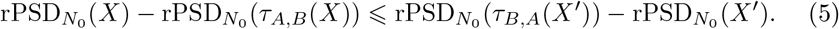

Note that *A, B* ⊆ *C*(*v*_0_), and therefore

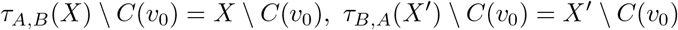

and thus 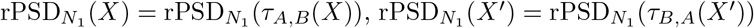. Assume first that *B*≠ ∅, so that in particular *X*^*′*^ ∩ *C*(*v*_0_)≠ ∅ and *τ*_*A,B*_(*X*) ∩ *C*(*v*_0_) ∅. Then,

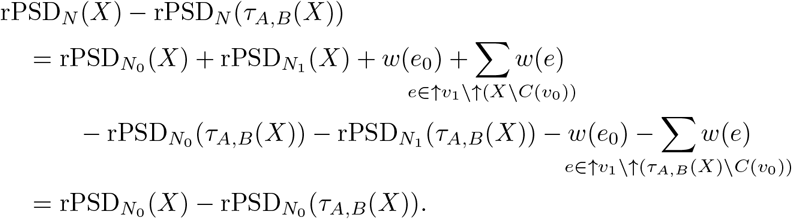

On the other hand we also have (by the same argument):

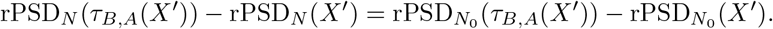

Therefore, by the induction hypothesis (Eqn. (5))

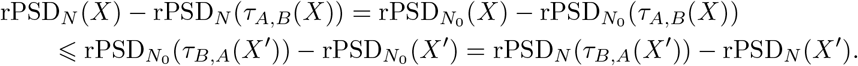

Assume now that *B* = ∅. Then, by the definition of *S*_*k,d*_, *A* must be a singleton and then *τ*_*A,B*_(*X*) ∩ *C*(*v*_0_) = ∅, because, by assumption, |*X* ∩ *C*(*v*_0_) | *>* 1. Then, arguing as above, we have that

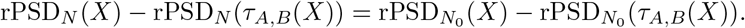

Similarly, if *X*^*′*^ ∩ *C*(*v*_0_)≠ ∅,

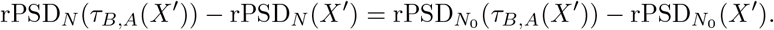

Otherwise, if *X*^*′*^ ∩ *C*(*v*_0_) = ∅,

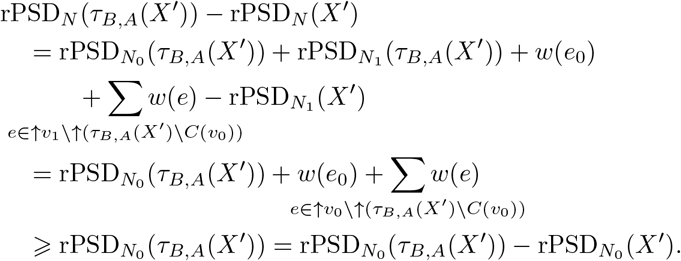

In either case applying the induction hypothesis (Eqn. (5)), we have again

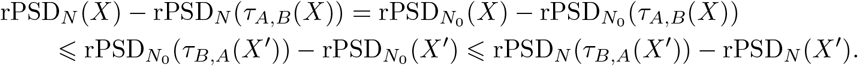
b. Assume now that the only node *v* in *T* such that |*X* ∩ *C*(*v*)|*>* |*X*^*′*^∩*C*(*v*)| and |*X* ∩*C*(*v*) | *>* 1 is the root *r*, and that *r* is not the split node of any blob in *N*, so that its children in *N* are also its children in *T*. Then, for every child *v* of *r*, if |*X* ∩ *C*(*v*) | *>* |*X*^*′*^∩ *C*(*v*) |, then |*X* ∩ *C*(*v*) | = 1. But since |*X*|*>* |*X*^*′*^|, *r* must have some child *v*_1_ such that |*X* ∩ *C*(*v*_1_) | *>* |*X*^*′*^ ∩ *C*(*v*_1_) | and hence such that *X*∩*C*(*v*_1_) = {*x*} and *X*^*′*^∩*C*(*v*_1_) = ∅; moreover, since |*X*| ⩾ 2, *r* has another child *v*_2_. For each *i* = 1, 2, let *N*_*i*_ be the subnetwork of *N* rooted at *v*_*i*_. Their sets of leaves *C*(*v*_*i*_) are pairwise disjoint and therefore, for each *Z* ⊆ Σ,

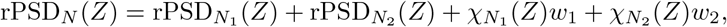

where each *w*_*i*_ is the weight of the edge (*r, v*_*i*_) and 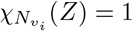 if *Z* ∩ *C*(*v*_*i*_) = ∅ and it is 0 otherwise. Take *A* = *{x}* and *B* = ∅. Then (*A, B*) ∈ 𝒮_0_(Σ), *τ*_*A,B*_(*X*) ∩ *C*(*v*_1_) = ∅, *τ*_*B,A*_(*X*^*′*^) ∩ *C*(*v*_1_) = *{x}, τ*_*A,B*_(*X*) ∩ *C*(*v*_2_) = *X* ∩ *C*(*v*_2_), and *τ*_*B,A*_(*X*^*′*^) ∩ *C*(*v*_2_) = *X*^*′*^ ∩ *C*(*v*_2_). Then:

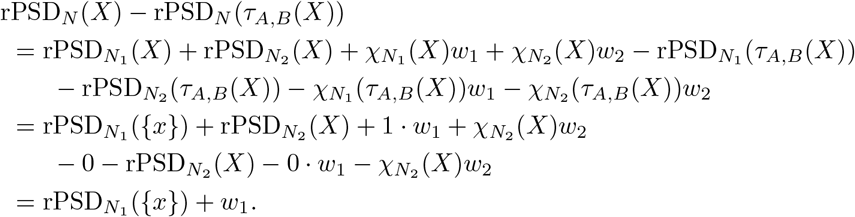

Similarly,

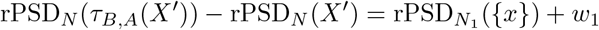

and hence, in this case,

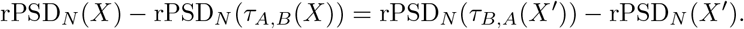
c. Assume now that the only node *v* in *T* such that |*X* ∩ *C*(*v*) |*>* |*X*^*′*^∩ *C*(*v*) | and |*X* ∩ *C*(*v*) | *>* 1 is the root *r*, and that it is the split node of a (single) blob. Furthermore, assume that there exists some exit reticulation *H* of that blob such that *X*^*′*^ ∩ *C*(*H*) = *X* ∩ *C*(*H*) = ∅. If *v*_1_, …, *v*_*d*′_ are the parents of *H*, with *d*^*′*^ ⩽ *d*, then let 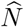 be the phylogenetic network obtained from *N* by removing the subnetwork *N*_*H*_, adding new leaves *h*_1_, …, *h*_*d*′_ with dummy labels outside Σ, and replacing each edge (*v*_*i*_, *H*) by an edge (*v*_*i*_, *h*_*i*_) with weight 0. 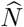 is still at-most-bifurcating, semi-*d*-ary, and level-*k* and it has less than *α* edges (we have removed the edges in *N*_*H*_). Therefore, by the induction hypothesis, it satisfies the thesis in the statement. Let 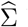 be its set of labels. Then, since, by assumption, 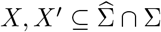, there exist *A* ⊆ *X \ X*^*′*^ and *B* ⊆ *X*^*′*^ *\ X* such that 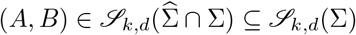,

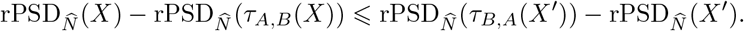

Since 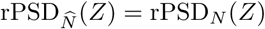 for every 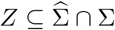, we conclude that

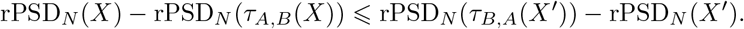
d. Finally, assume that the root is the only node *v* in *T* such that |*X* ∩ *C*(*v*)| *>* |*X*^*′*^ ∩ *C*(*v*)| and |*X* ∩ *C*(*v*)| *>* 1, and that it is the split node of a blob ℬ with no exit reticulation *H* having *X*^*′*^ ∩ *C*(*H*) = *X* ∩ *C*(*H*) = ∅.

Let ℬ^∗^ be the set of nodes of ℬ that have a child outside of ℬ; if *v* ∈ *ℬ*^∗^, we shall denote its child outside of ℬ by 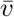. Notice that the exit reticulations of ℬ belong to ℬ^∗^, that *r* ∉ ℬ^∗^ (its two children must belong to the blob), and that, since reticulations have out-degree 1, the internal reticulations of ℬ do not belong to ℬ^∗^. For each *v* ∈ *ℬ*^∗^ let 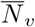 be the subnetwork of *N* rooted at *v* consisting of 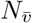, *v* and the edge 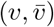.

For each *Z* ⊆ Σ, consider the multiset 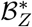 of nodes of ℬ^∗^ with support

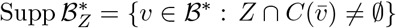

and multiplicities 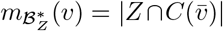. Since the subnetworks 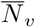, with *v* ∈ ℬ^∗^, have pairwise disjoint sets of leaves and the union of their sets of leaves is Σ, we have that 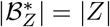 and

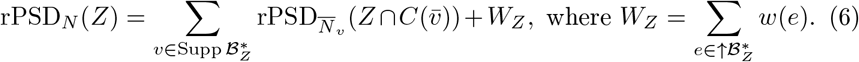

So,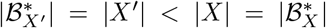; by the current assumption, every exit reticulation belongs to 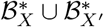; and if 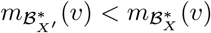, then 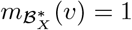. Therefore, the multisets 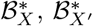; satisfy the hypotheses of Lemma 5 and thus there exist a set ℬ_*A*_ and a multiset ℬ_*B*_ of nodes of ℬ such that:

1. ℬ_*B*_ = ∅ and |ℬ_*A*_| = 1 or

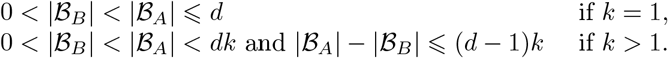
2. ℬ_*A*_ ⊆ Supp 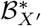 \ Supp 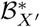. Therefore, if 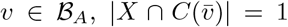 and 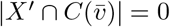.
3. 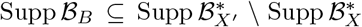. and, for every *v* ∈ Supp ℬ_*B*_, 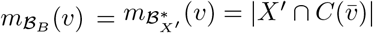.
4. 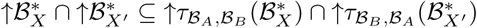

Let

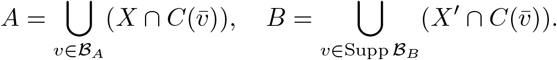

Then, *A* ⊆ *X \X*^*′*^, *B* ⊆ *X*^*′*^ *\X*, 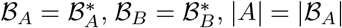, and |*B*| = |*ℬ*_*B*_|. In particular, by property (1), (*A, B*) ∈ 𝒮_*k,d*_(Σ). Moreover

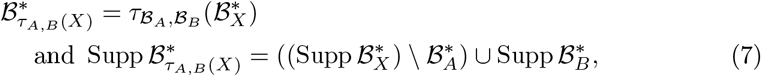

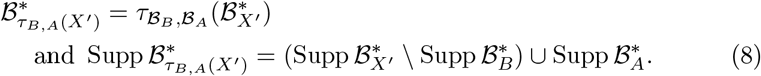

Indeed, as to Eqn. (7),

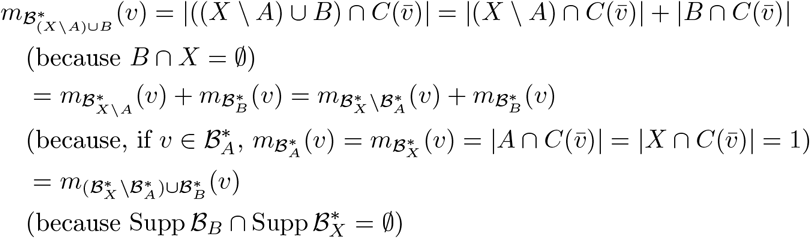

and then, again because 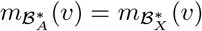 for every 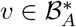,

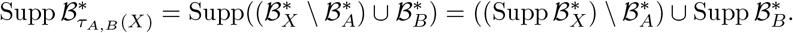

A similar argument, using that, for every 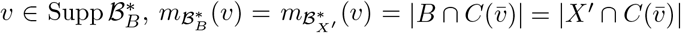 and that Supp *B*_*A*_ ∩ Supp 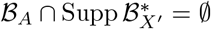, proves Eqn. (8).

We want to prove that

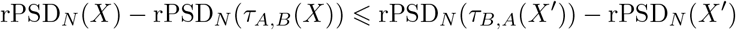

Now, by Eqn. (6),

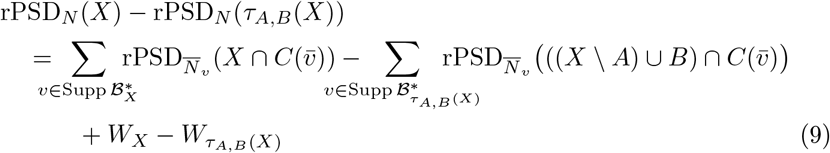

where

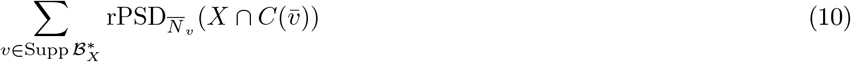

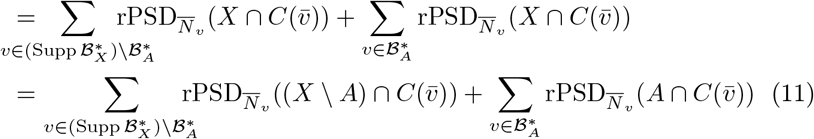

because if 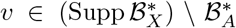, then 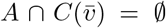, and if 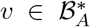, then 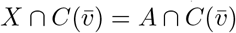; and

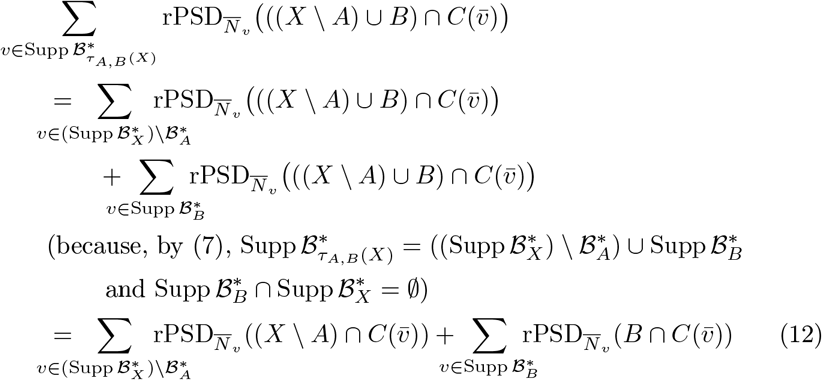

because if 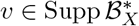, then 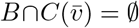, and if 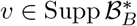, then 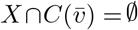.

Therefore, combining Eqns. (9) to (12), we obtain

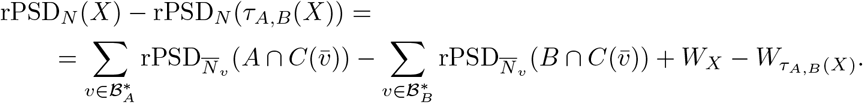

A similar argument, using (8) instead of (7), proves that

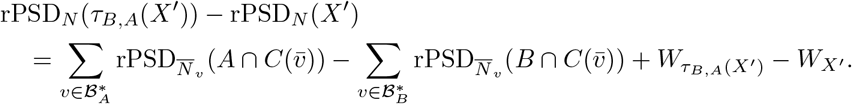

Thus,

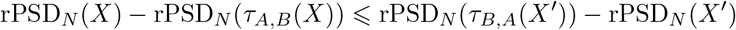

if, and only if,

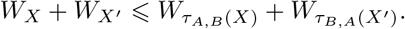

Finally, this last inequality holds because

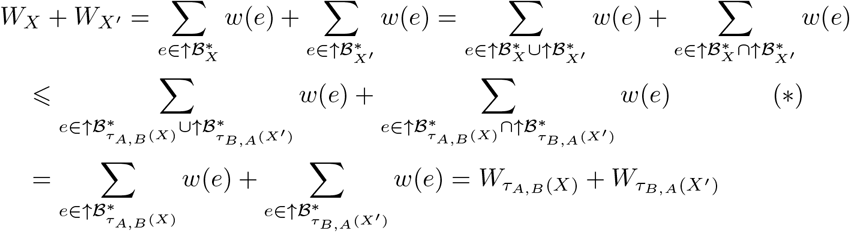

where step (∗) is due to

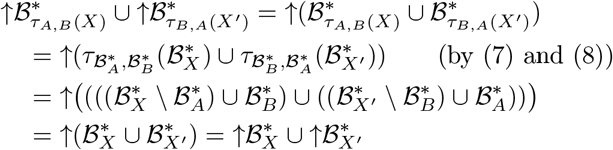

and, by property (4) of ℬ_*A*_ and ℬ_*B*_ (and, again, (7) and (8)),

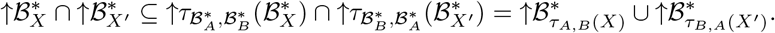

This completes the proof of case **(d)**.

There are semi-*d*-ary level-*k* weighted phylogenetic networks *N* on sets of taxa Σ and pairs of sets of leaves *X, X*^*′*^⊆ Σ with |*X*^*′*^| *<* |*X*| having no rPSD_*N*_ - improving pair (*A, B*) with |*A*|− |*B*| *<* (*d* − 1)*k*. So, in this sense, the set 𝒮_*k,d*_(Σ) cannot be improved. The next example describes one such network for *d* = 2; it is straightforward to generalize it to the semi-*d*-ary setting for any *d* ⩾ 2

### Example 7.

Consider the semibinary level-*k* phylogenetic network *N* with set of nodes and edges

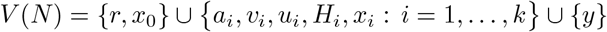

and edges

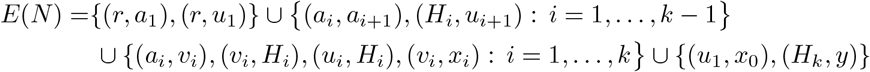

and set of labels (which we indentify with its leaves) Σ = {*y, x*_1_, …, *x*_*k*_} ; cf. Fig. 3. Assume that all its edges have weight *>* 0. Given an edge *e* = (*s, t*), we denote its weight by *w*(*e*) or *w*(*s, t*).

**Figure 3:**
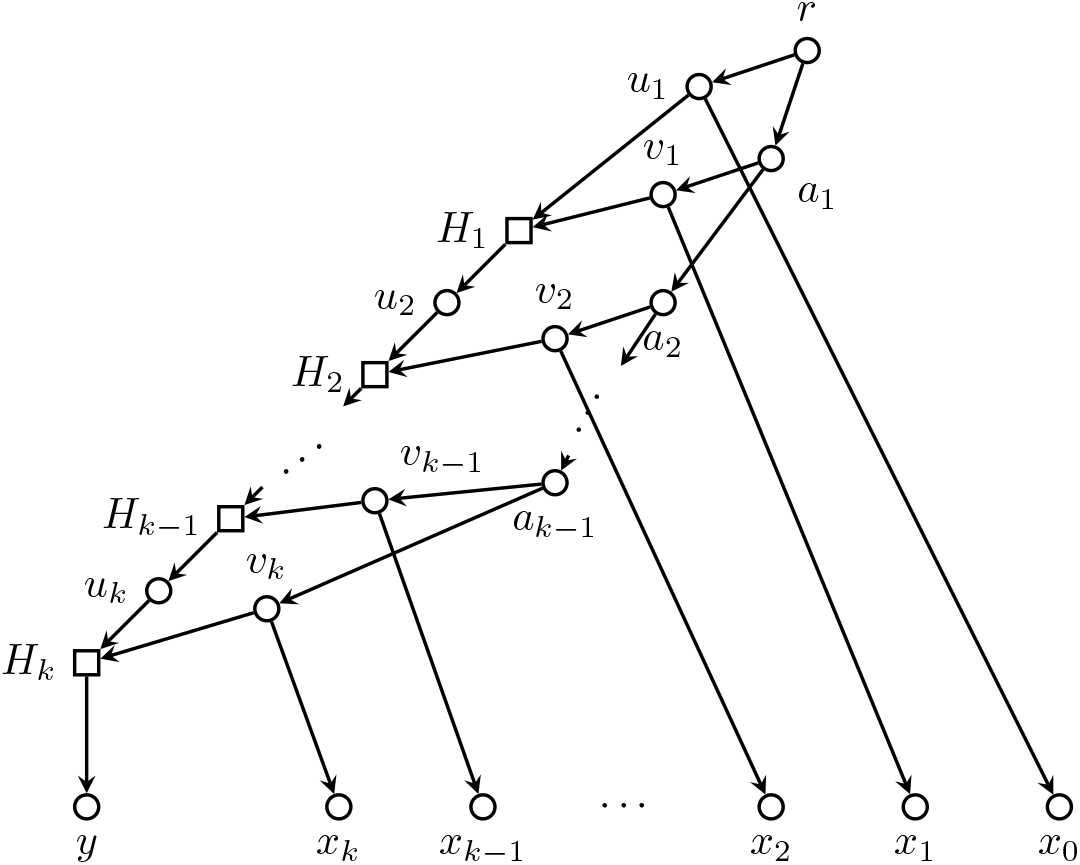
The network *N* in Example 7.

Let *X* = {*x*_0_, *x*_1_, …, *x*_*k*_} and *X*^*′*^ = {*y*}. Let us check that, for every (*A, B*) such that *A* ⊆ *X, B* ⊆ *X*^*′*^ and |*B*| *<* |*A*|

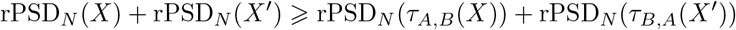

and that the equality holds only when (*A, B*) = (*X, X*^*′*^). This will imply that the only rPSD_*N*_ -improving pair for *X, X*^*′*^ in 𝒮_*k*,2_(Σ) is (*X, X*^*′*^).

Let *E*_0_ be the edges in ↑{*u*_1_, *v*_1_, …, *v*_*k*_} and let *E*_1_ = *E*(*N*) \ (*E*_0_∪{(*v*_*i*_, *x*_*i*_)}_*i*=1,…,*k*_). Then,

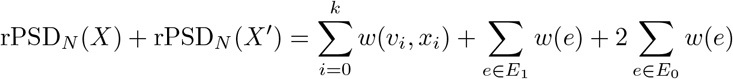

Now, on the one hand, if *B* = ∅ and *A*≠= ∅

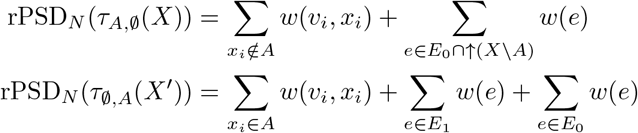

and then

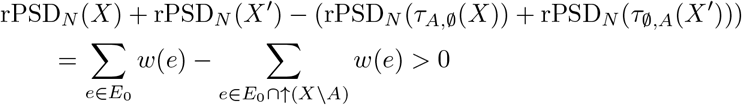

because *X* \ *A* ≠*X* and for every *x*_*i*_ ∈ *X* the edge (*a*_*i*_, *v*_*i*_) (or (*r, u*_1_) if *i* = 0) does not belong to ↑(*X* \ {*x*_*i*_}).

And, on the other hand, if *B* = {*y*},

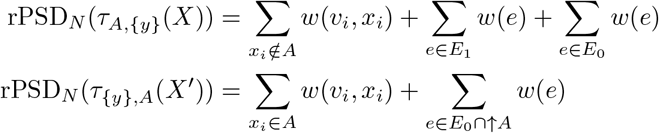

and

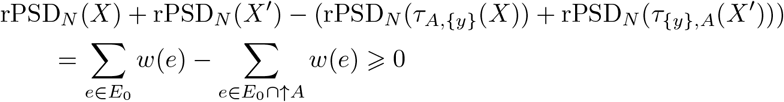

and, arguing as above, the inequality is an equality exactly when *A* = *X*.

### Example 8.

Example 7 shows that there are networks and sets *X, X*^*′*^ such that any rPSD-improving pair (*A, B*) must have |*A*| − |*B*| = (*d* − 1)*k*. We now show that this is true even for the special case where |*X*| = |*X*^*′*^|− 1 (which is common in Section 5). Consider the weighted semi-3-ary level-1 phylogenetic network depicted in Figure 4. In it, *X* = {*x*_00_, *x*_01_, *x*_02_, *z*_1_} and *X*^*′*^ = {*x*_11_, *x*_12_, *z*_0_} do not have any rPSD-improving pair (*A, B*) in 𝒮_1,3_ = 𝒮_2,2_ with |*A*| − |*B*| = 1.

**Figure 4:**
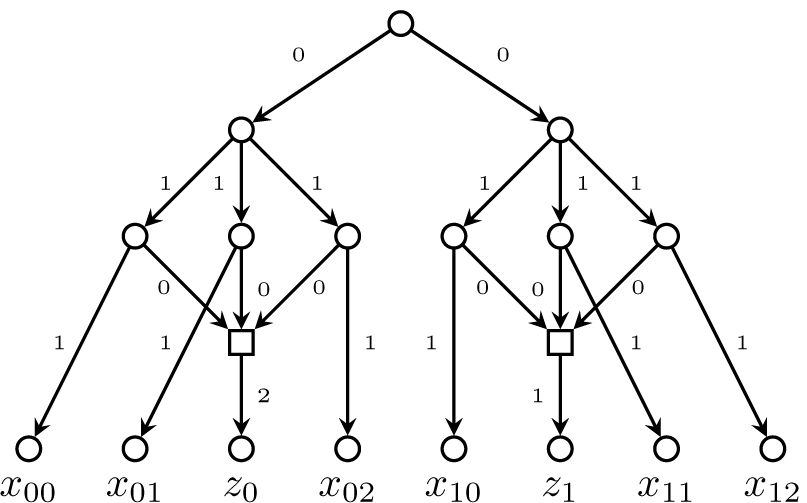
The network in Example 8.

We close this section with a refinement of Theorem 6 for semi-*d*-ary level-1 networks. The proof is similar to that of the aforementioned theorem, and we postpone it until an appendix at the end of the paper.

### Corollary 9.

*If N is a semi-d-ary level-1 weighted phylogenetic network on* Σ, rPSD_*N*_ *satisfies the exchange property with respect to*

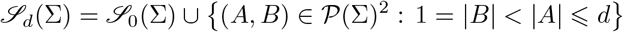

*Moreover, if X, X*^*′*^ *have an improving pair* (*A, B*) ∈ 𝒮_*d*_(Σ) \ 𝒮_0_(Σ), *then there exists a blob in N with exit reticulation H and split node v such that X* ∩ *C*_*N*_ (*H*) = ∅, *B* ⊆ *C*_*N*_ (*H*), *and A* ⊆ *C*_*N*_ (*v*).

## 5 Applications

In this section we apply Theorem 6 to the study of rPSD_*N*_ -optimal subsets for low values of the level of *N* and of the in-degree of its reticulations. Throughout this section, let *N* be a weighted phylogenetic network on a finite set Σ of cardinality *n*. We shall use the following notations:

- For every *m* = 0, …, *n*, let

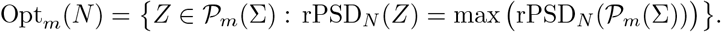

By convention, if *m <* 0 or if *m > n*, we set Opt_*m*_(*N*) = ∅.
- For every *k* ⩾ 1 and *d* ⩾ 2, for every 1 ⩽ *j* ⩽ (*d* − 1)*k*, and for every *X* ∈ 𝒫 (Σ), let

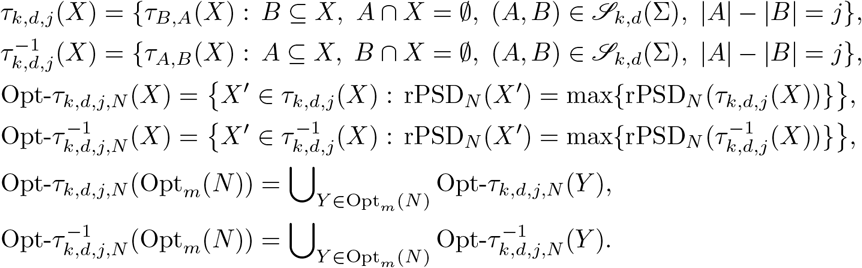

Notice that *X*^*′*^ ∈ *τ*_*k,d,j*_(*X*) if, and only if, 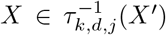, and that if *X*^*′*^ ∈ *τ*_*k,d,j*_(*X*), then |*X*^*′*^| = |*X*| + *j*.

When *N* is clear from the context, we shall simply write rPSD, Opt_*m*_, Opt-*τ*_*k,d,j*_(*X*), and 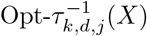. Now we define a relation to be used in this section together with basic results about it.

### Definition 10.

An *optimal sequence* of *N* is a sequence *Y* = (*Y*_*p*_)_0⩽*p*⩽*n*_ where each *Y*_*p*_ ∈ Opt_*p*_(*N*).

### Definition 11.

Let *N* be a weighted semi-*d*-ary level-*k* phylogenetic network on Σ and *Y* an optimal sequence of *N*. For each 0 ⩽ *p, q, p*^*′*^, *q*^*′*^ ⩽ *n*, define 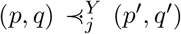 to be true if, and only if, *p > q* and there exists a rPSD_*N*_ - improving pair (*A, B*) ∈ 𝒮_*k,d*_(Σ) for *Y*_*p*_, *Y*_*q*_ such that |*A*| − |*B*| = *j, p*^*′*^ = *p* − *j*, and *q*^*′*^ = *q* + *j*. When we need to emphasize the rPSD_*N*_ -improving pair of sets (*A, B*), we shall say that “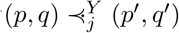 by the improving pair (*A, B*)”.

In addition, we shall use the following shorthand notation {-, -}:

- on the right: 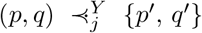 is equivalent to 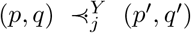 or 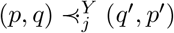,
- on the left: 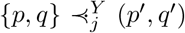 is equivalent to (max{*p, q*}, min 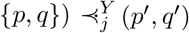.

### Remark 12.

By Theorem 6, given any optimal sequence *Y* of a semi-*d*-ary level-*k* phylogenetic network and *Y*_*p*_, *Y*_*q*_ in *Y* with 0 ⩽ *q < p*, we have 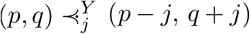 for some 1 ⩽ *j* ⩽ (*d* − 1)*k*.

### Lemma 13.

*Let N be a weighted phylogenetic network on* Σ *and Y an optimal sequence of N*. *If* 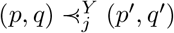 *by the improving pair* (*A, B*) *and* rPSD(*Y*_*p′*_)+ rPSD(*Y*_*q′*_) ⩽ rPSD(*Y*_*p*_) + rPSD(*Y*_*q*_), *then*

- *τ*_*A,B*_(*Y*_*p*_) ∈ Opt_*p′*_ *and hence Y*_*p*_ ∈ Opt*-τ*_*k,d,j*_(Opt_*p′*_),
- *τ*_*B,A*_(*Y*_*q*_) ∈ Opt_*q′*_ *and hence* 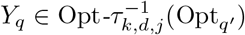.

*Proof*. If 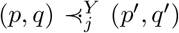 by the improving pair (*A, B*), the optimality of *Y*_*p′*_ and *Y*_*q*′_ implies that

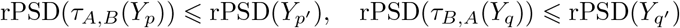

and then, if rPSD(*Y*_*p′*_) + rPSD(*Y*_*q′*_) ⩽ rPSD(*Y*_*p*_) + rPSD(*Y*_*q*_),

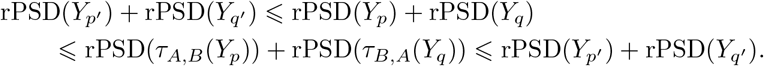

We deduce that

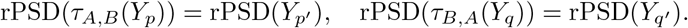

Therefore

- *τ*_*A,B*_(*Y*_*p*_) ∈ Opt_*p′*_ and thus *Y*_*p*_ = *τ*_*B,A*_(*τ*_*A,B*_(*Y*_*p*_)) ∈ *τ*_*k,d,j*_(Opt_*p′*_); actually, being *Y*_*p*_ optimal, *Y*_*p*_ ∈ Opt-*τ*_*k,d,j*_(Opt_*p′*_),
- *τ*_*B,A*_(*Y*_*q*_) ∈ Opt_*q′*_ and thus 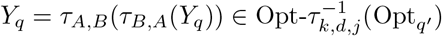.

### Corollary 14.

*Let N be a weighted phylogenetic network on* Σ *and Y an optimal sequence of N*. *If there exists a closed* 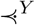*-chain of length m* ⩾ 1

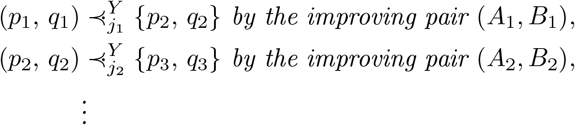

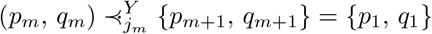 *by the improving pair* (*A*_*m*_, *B*_*m*_),

*then, for each* 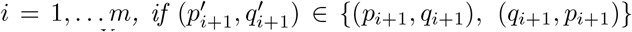 *is the pair satisfying* 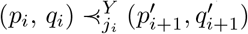,

- 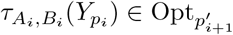 *and* 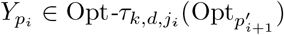,
- 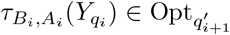 *and* 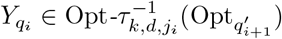,

*Proof*. The closed chain ensures that all the inequalities in

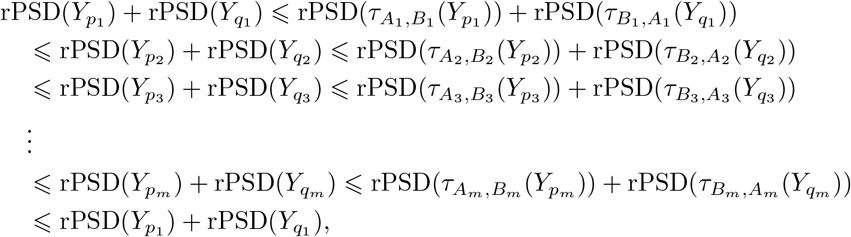

are equalities, and the result follows from applying the Lemma 13 at each 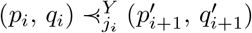.

M. Bordewich *et al* proved in [4, Cor 4.6] that the optimization problem for rPSD_*N*_ can be solved in polynomial time when *N* is a galled tree (a semibinary level-1 network). The next result strengthens their result by providing a recursive construction of the rPSD_*N*_ -optimal sets for these networks.

### Proposition 15.

*If N is a weighted galled tree, for every m* = 1, …, *n*

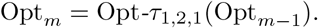

*Proof*. Let *Y* be an optimal sequence of *N* and fix 1 ⩽ *m* ⩽ *n*. Then, by Theorem 6,

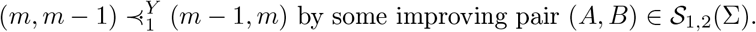

Thus, by Corollary 14, we have

- *τ*_*A,B*_(*Y*_*m*_) ∈ Opt_*m*−1_ and hence *Y*_*m*_ ∈ Opt-*τ*_1,2,1_(Opt_*m*−1_),
- *τ*_*B,A*_(*Y*_*m*−1_) ∈ Opt_*m*_ and hence Opt-*τ*_1,2,1_(*Y*_*m*−1_) ⊆ Opt_*m*_.

Since the choice of *Y* is arbitrary, we conclude that

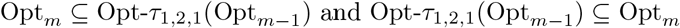

as stated.

### Remark 16.

Notice that along the proof of the previous proposition we have proved that, in a weighted galled tree, for every *Y*_*m*_ ∈ Opt_*m*_ and *Y*_*m*−1_ ∈ Opt_*m*−1_, there exists some pair (*A, B*) ∈ 𝒮_1,2_(Σ), with *A* ⊆ *Y*_*m*_ *\ Y*_*m*−1_ and *B* ⊆ *Y*_*m*−1_ *\ Y*_*m*_, such that *τ*_*A,B*_(*Y*_*m*_) ∈ Opt_*m*−1_ and *τ*_*B,A*_(*Y*_*m*−1_) ∈ Opt_*m*_.

So, on weighted galled trees and for every *k* = 1, …, *n*, all members of Opt_*k*_ are obtained from members of Opt_*k* − 1_ by either optimally adding a leaf or optimally replacing a leaf by a pair of leaves. This yields a simple greedy polynomial time algorithm to compute the family of optimal sets Opt_*k*_ (*N*) in increasing order of *k* that extends the greedy Algorithm 1 for phylogenetic trees:

### Algorithm 2: Greedy forward for galled trees

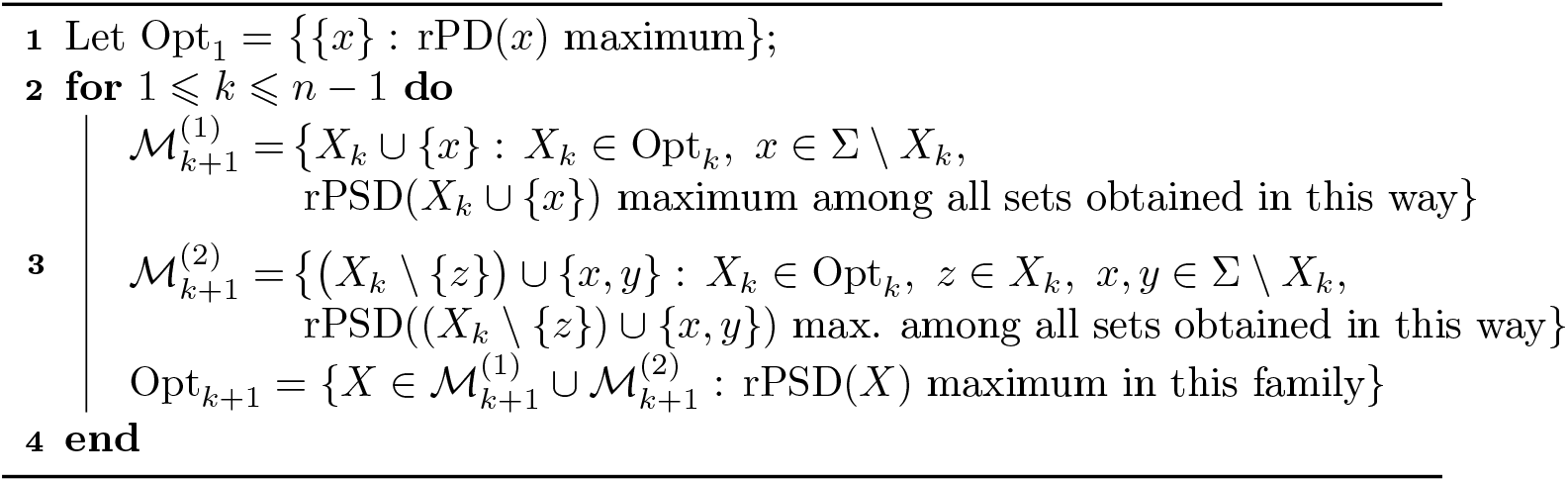

Moreover, Proposition 15 also entails that, on weighted galled trees and for every *k* = 1, …, *n* all members of Opt_*k* − 1_ are obtained from members of Opt_*k*_ by removing a leaf or replacing a pair of leaves by a leaf in such a way that the value of rPSD decreases the least. This yields a similar backwards greedy algorithm that recurrently computes Opt_*k*−1_ from Opt_*k*_ (Algorithm 3).

Let us move now one step up in the complexity ladder of phylogenetic networks. Before proceeding, notice that

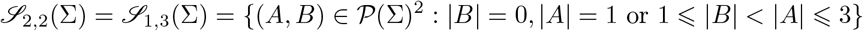

and in particular, for every *j* = 1, 2, Opt-*τ*_1,3,*j*_ = Opt-*τ*_2,2,*j*_.

### Proposition 17.

*If N is a weighted semi-binary level-2 or a semi-3-ary level-1 network on* Σ, *then:*

a. Opt_*m*_ ⊆ Opt*-τ*_2,2,1_(Opt_*m*−1_) ∪ Opt*-τ*_2,2,2_(Opt_*m*−2_) *for every m* = 1, …, *n*.
b. 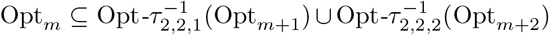*for every m* = 1, …, *n* − 1. *Proof*. To simplify the notations, we shall denote Opt-*τ*_1,3,*j*_ = Opt-*τ*_2,2,*j*_ by simply Opt-*τ*_*j*_.

### Algorithm 3: Greedy backward for galled trees

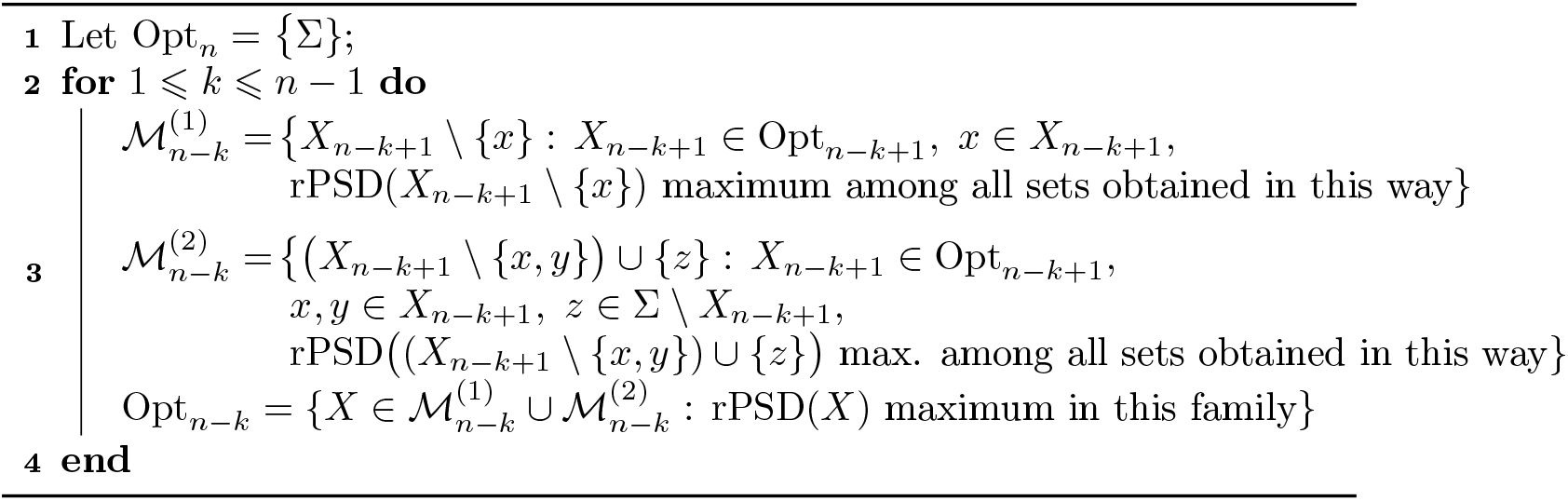

Let *Y* be an optimal sequence of *N* and fix 1 ⩽ *m* ⩽ *n*. Then, by Theorem 6,

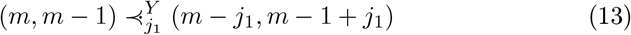

for some *j*_1_ = 1 or *j*_1_ = 2.

1. If *j*_1_ = 1, then 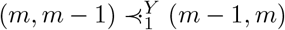, and hence, as in Proposition 15,

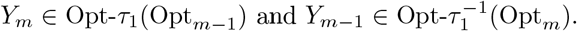
2. If *j*_1_ = 2, then 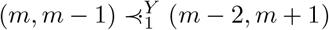. Applying Theorem 6 again,

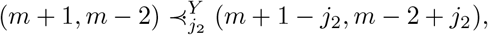

for some *j*_2_ = 1 or *j*_2_ = 2. In both cases, {*m*+1−*j*_2_, *m*−2+*j*_2_} = {*m*−1, *m*}, thus closing the 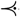-chain initiated with (13). Then, by Corollary 14,

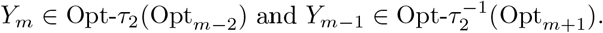

Note that in both cases we have
  - *Y*_*m*_ ∈ Opt-*τ*_1_(Opt_*m*−1_) ∪ Opt-*τ*_2_(Opt_*m*−2_),
  - 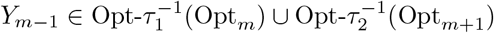, which, by the arbitrary choice of *Y* and *m*, concludes the proof.

### Algorithm 4: Greedy forward for semibinary level-2 or semi-3-ary level-1 networks

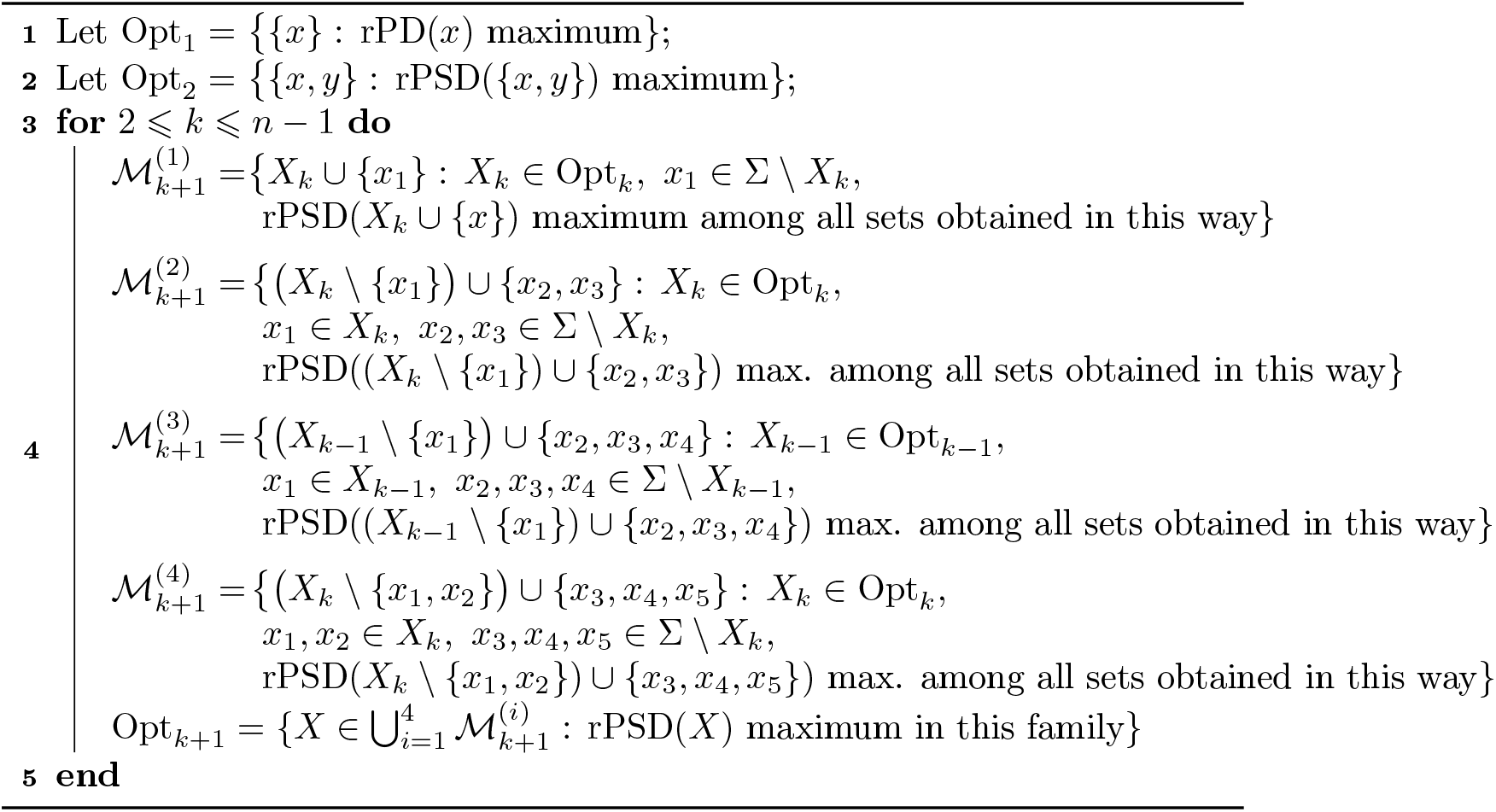

So, point (a) in the last proposition tells us that if *N* is semibinary level-2 or semi-3-ary level-1, all members of Opt_*k*_ are obtained either from members of Opt_*k* − 1_ by optimally adding a leaf, or optimally replacing a leaf by a pair of leaves, or optimally replacing a pair of leaves by a triple of leaves (this possibility need not be considered in the semi-3-ary level-1 case by Corollary 9), or from members of Opt_*k* − 2_ by optimally replacing a leaf by a triple of leaves. This proves the correctness of the following polynomial time greedy algorithm to compute the family of optimal sets Opt_*k*_ for such a network *N* in increasing order of *k* (as we have mentioned, if *N* is semi-3-ary level-1, the sets ℳ^(4)^ in the loop need not be computed).

Point (b) in the last proposition yields a similar greedy algorithm that computes the optimal sets Opt_*k*_ for such a network in *decreasing* order of *k*. We leave the details to the reader.

### Example 18.

Consider the weighted phylogenetic networks in Figure 5: On the left, a semi-3-ary level-1 network (with the same topology as Example 8, but different weights) and on the right right a semibinary level-2 network obtained by blowing up the reticulations in the left-hand side network into a pair of in-degree 2 connected reticulations.

**Figure 5:**
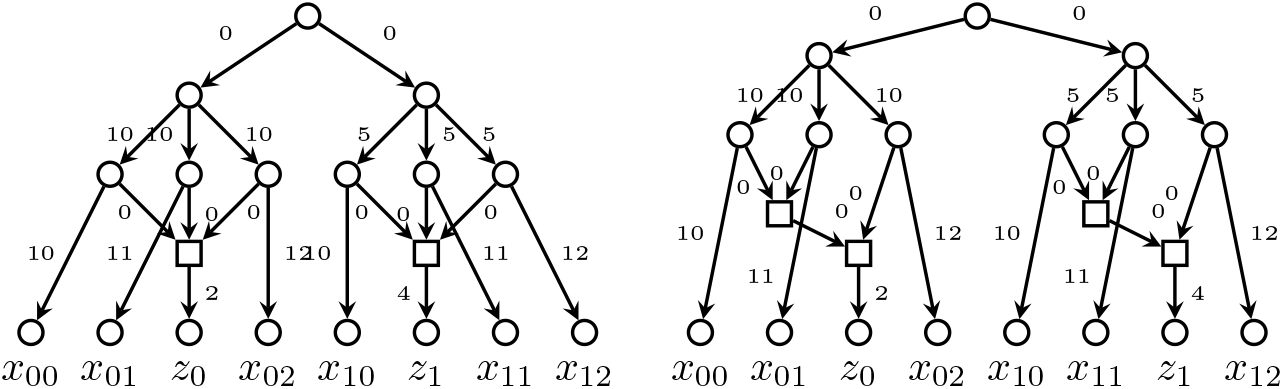
Networks from Example 18.

We have the following optimal sets of leaves:

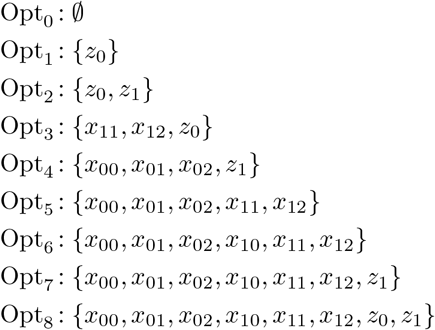

In both cases, {*x*_00_, *x*_01_, *x*_02_, *z*_1_} ∈ Opt_4_ \ Opt-*τ*_1_(Opt_3_) and 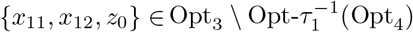.

Now, if we move one more step in the complexity ladder, the structure of the optimal sets is no longer so simple.

### Proposition 19.

*If N is a weighted semi-binary level-3 or a semi-4-ary level-1 network on* Σ, *then, for every m* = 1, …, *n, at least one of the following assertions is true:*

- Opt_*m*+1_ = Opt*-τ*_*k,d*,3_(Opt_*m*−2_),
- 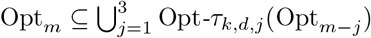 *and* 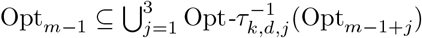.

*(where* (*k, d*) *is* (3, 2) *or* (1, 4), *depending on the type of network)*.

*Proof*. To begin with, notice that

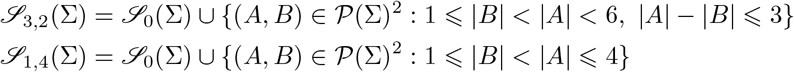

and therefore 𝒮_1,4_(Σ) ⊆ 𝒮_3,2_(Σ). To simplify the notations, we shall abbreviate Opt-*τ*_*k,d,j*_ by simply Opt-*τ*_*j*_. Observe that in both cases considered in the statement *j* can go from 1 to 3.

Let *Y* be an optimal sequence of *N* and fix 1 *< m* ⩽ *n*. Then, by Theorem 6,

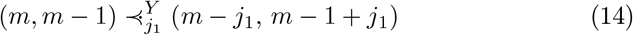

for some *j*_1_ ∈ {1, 2, 3}.

1. If *j*_1_ = 1, then, we conclude as in (1) in the proof of Proposition 17 that *Y*_*m*_ ∈ Opt-*τ*_1_(Opt_*m*−1_) and 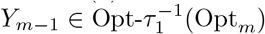.
2. If *j*_1_ = 2, then (*m* − *j*_1_, *m* − 1 + *j*_1_) = (*m* − 2, *m* + 1). Applying Theorem 6 again,

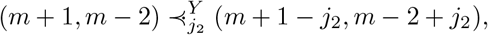

for some *j*_2_ ∈ {1, 2, 3}.
  (2.a) If *j*_2_ = 1 or *j*_2_ = 2, we conclude as in (2) in the proof of Proposition 17 that *Y*_*m*_ ∈ Opt-*τ*_2_(Opt_*m*−2_) and 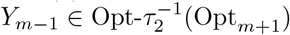.
  (2.b) When *j*_2_ = 3, we have 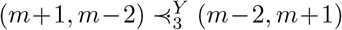 and we can only deduce that *Y*_*m*+1_ ∈ Opt-*τ*_3_(Opt_*m*−2_) and 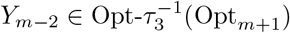.
3. If *j*_1_ = 3, then (*m −j*_1_, *m* −1 + *j*_1_) = (*m* −3, *m* + 2). Applying Theorem 6 again,

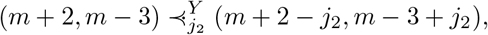

for some *j*_2_ ∈ {1, 2, 3}.
  (3.a) If *j*_2_ = 1, then 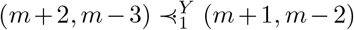. Applying Theorem 6, we have

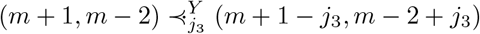

for some *j*_3_ ∈ {1, 2, 3}.
    (3.a.i) If *j*_3_ = 1 or *j*_3_ = 2, then 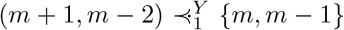, closing the 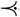-chain initiated with (14). Then, by Corollary 14, *Y*_*m*_ ∈ Opt-*τ*_3_(Opt_*m*−3_) and 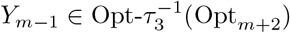.
    (3.a.ii) If *j*_3_ = 3, then 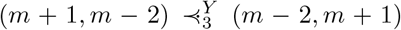 as in (2.b) and we only have that *Y*_*m*+1_ ∈ Opt-*τ*_3_(Opt_*m*−2_) and 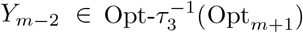.
  (3.b) If *j*_2_ = 2 or *j*_2_ = 3, then 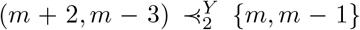, closing the 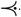-chain initiated with (14). and then, by Corollary 14, *Y*_*m*_ ∈ Opt-*τ*_3_(Opt_*m*−3_) and 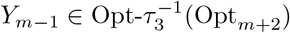.

Summarizing, we only have two possibilities:

- On the one hand, in the cases (1), (2.a), (3.a.i), and (3.b),

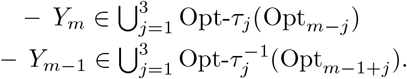
- On the other hand, in the cases (2.b) and (3.a.ii),

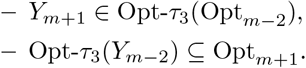

By the arbitrary choice of *Y* and *m*, this concludes the proof.

A similar result holds for (*k, d*) such that (*d* − 1)*k* = 4.

### Proposition 20.

*If N is a weighted semi-5-ary level-1 or a semi-3-ary level-2 network on* Σ, *then, for every m* = 1, …, *n, at least one of the following assertions is true:*

- Opt-_*m+1*_ *=* Opt-*τ*_*k,d,3*_ (Opt_*m+2*_)
- 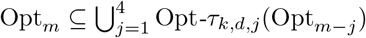 *and* 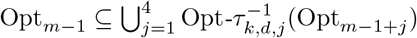

*(where k, d are* 2, 3 *or* 1, 5, *depending on the type of network)*.

*Proof*. To begin with, notice that

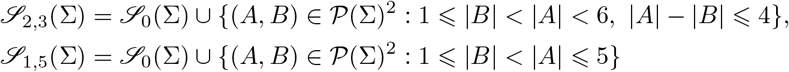

and therefore 𝒮 _1,5_(Σ) = 𝒮 _2,3_(Σ). To simplify the notations, we shall abbreviate Opt-*τ*_1,5,*j*_ = Opt-*τ*_2,3,*j*_ by simply Opt-*τ*_*j*_. Observe that in both cases considered in the statement *j* can go from 1 to 4.

Let *Y* be an optimal sequence of *N* and fix 1 *< m* ⩽ *n*. Then, by Theorem 6,

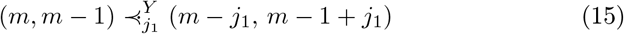

for some *j*_1_ ∈ {1, 2, 3, 4}.

1. If *j*_1_ = 1, then, we conclude as in (1) in the proof of Proposition 17 that *Y*_*m*_ ∈ Opt-*τ*_1_(Opt_*m*−1_) and 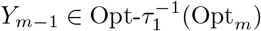.
2. If *j*_1_ = 2, then (*m − j*_1_, *m −* 1 + *j*_1_) = (*m −* 2, *m* + 1). Applying Theorem 6 again,

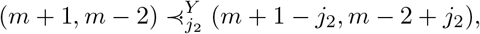

for some *j*_2_ ∈ {1, 2, 3, 4}.
  (2.a) If *j*_2_ = 1 or *j*_2_ = 2, we conclude as in (2) in the proof of Proposition 17 that *Y*_*m*_ ∈ Opt-*τ*_2_(Opt_*m*−2_) and 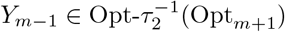.
  (2.b) When *j*_2_ = 3, we have 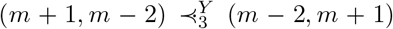 and, as in (2.b) in the proof of Proposition 17, we can only conclude that *Y*_*m*+1_ ∈ Opt-*τ*_3_(Opt_*m*−2_) and 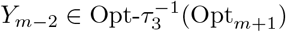.
  (2.c) When *j*_2_ = 4, we have 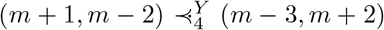. Applying Theorem 6 again,

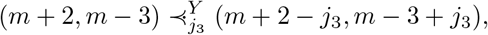

for some *j*_3_ ∈ {1, 2, 3, 4}. Now:
    (2.c.i) If *j*_3_ = 1 or 4, {*m* + 2 − *j*_3_, *m* − 3 + *j*_3_} = {*m* + 1, *m* − 2} and, as in (2.b), we conclude that *Y*_*m*+1_ ∈ Opt-*τ*_3_(Opt_*m*−2_) and 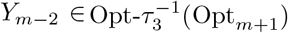.
    (2.c.ii) If *j*_3_ = 2 or 3, {*m* + 2 − *j*_3_, *m* − 3 + *j*_3_} = {*m, m* − 1} and, as in (2.a), we conclude that *Y*_*m*_ ∈ Opt-*τ*_2_(Opt_*m*−2_) and 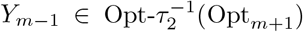.
3. If *j*_1_ = 3, then (*m − j*_1_, *m −* 1 + *j*_1_) = (*m −* 3, *m* + 2). Applying Theorem 6 again,

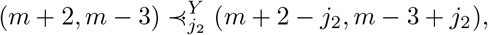

for some *j*_2_ ∈ {1, 2, 3, 4}.
  (3.a) If *j*_2_ = 2 or 3, {*m*+2−*j*_2_, *m*−3+*j*_2_} = {*m, m*−1}, closing the 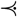-chain initiated with (15). Then, by Corollary 14, *Y*_*m*_ ∈ Opt-*τ*_3_(Opt_*m*−3_) and 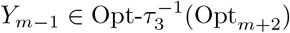.
  (3.b) If *j*_2_ = 1 or 4, {*m* + 2 − *j*_2_, *m −* 3 + *j*_2_} = {*m* + 1, *m −* 2}. But now we can follow as in case (2) and we conclude that one of the following situations must hold:
    - *Y*_*m*_ ∈ Opt-*τ*_3_(Opt_*m*−3_) and 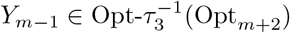,
    - *Y*_*m+1*_ ∈ Opt-*τ*_3_(Opt_*m*−2_) and 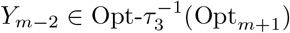.
4. If *j*_1_ = 4, then (*m − j*_1_, *m* − 1 + *j*_1_) = (*m* − 4, *m* + 3). Applying Theorem 6 again,

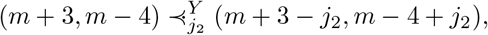

for some *j*_2_ ∈ {1, 2, 3, 4}.
  (4.a) If *j*_2_ = 3 or 4, {*m*+3−*j*_2_, *m*−4+*j*_2_} = {*m, m*−1}, closing the 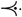-chain initiated with (15). Then, by Corollary 14, *Y*_*m*_ ∈ Opt-*τ*_4_(Opt_*m*−4_) and 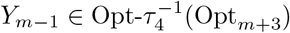.
  (4.b) If *j*_2_ = 1, 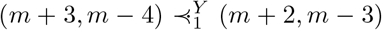 and we can follow as in (3), obtaining that one of the following situations must hold:
    - *Y*_*m*_ ∈ Opt-*τ*_4_(Opt_*m*−4_) and 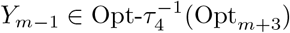,
    - *Y*_*m+1*_ ∈ Opt-*τ*_3_(Opt_*m*−2_) and 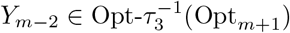.
  (4.c) If *j*_2_ = 2, 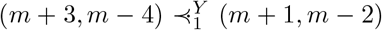 and we can follow as in (2), obtaining that one of the following situations must hold:
    - *Y*_*m*_ ∈ Opt-*τ*_4_(Opt_*m*−4_) and 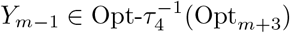,
    - *Y*_*m+1*_ ∈ Opt-*τ*_3_(Opt_*m*−2_) and 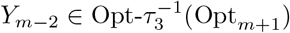.

Summarizing, we have two possibilities: either

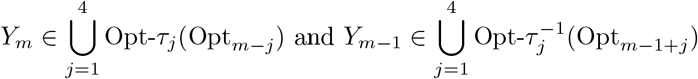

or

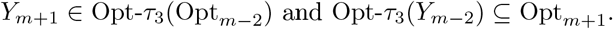

By the arbitrary choice of *Y* and *m*, this concludes the proof.

Now, while we could provide a greedy optimization algorithm for semibinary level-2 networks or semi-3-ary level-1 networks, an analogous argument fails for higher-level networks. The reason why Propositions 19 and 20 are not sufficient to provide such a greedy algorithm is that we would require their second statement —or a similar expression— to be true for all *m*. In the occurrence of any *m* where only the first statement in these proposition holds (this is, Opt_*m*+1_ = Opt-*τ*_*k,d*,3_(Opt_*m*−2_)), we do not have enough information about Opt_*m*_ to be able to ensure that it can be obtained from previous results. A similar problem occurs for the backward version of the algorithm.

But we must point out that, despite the last two propositions, we have not been able to find any weighted semi-binary level-3 or any semi-4-ary level-1 network on Σ for which 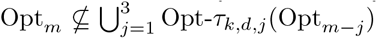 for some *m*. Similarly, we have not been able to find any weighted semi-5-ary level-1 or any semi-3-ary level-2 network for which 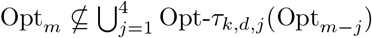 for some *m*. So, it might be possible that the greedy algorithm also works in these cases, since we have not discovered a counterexample that disproves its correctness for these types of networks.

We close this chapter with several examples that illustrate our search for a counterexample. More examples illustrating our search for a counterexample and the challenges encountered can be found in the second author’s PhD Thesis [30].

### Example 21.

Consider the weighted phylogenetic networks in Figure 6: On the left, a semi-4-ary level-1 network and on the right a semibinary level-3 network.

**Figure 6:**
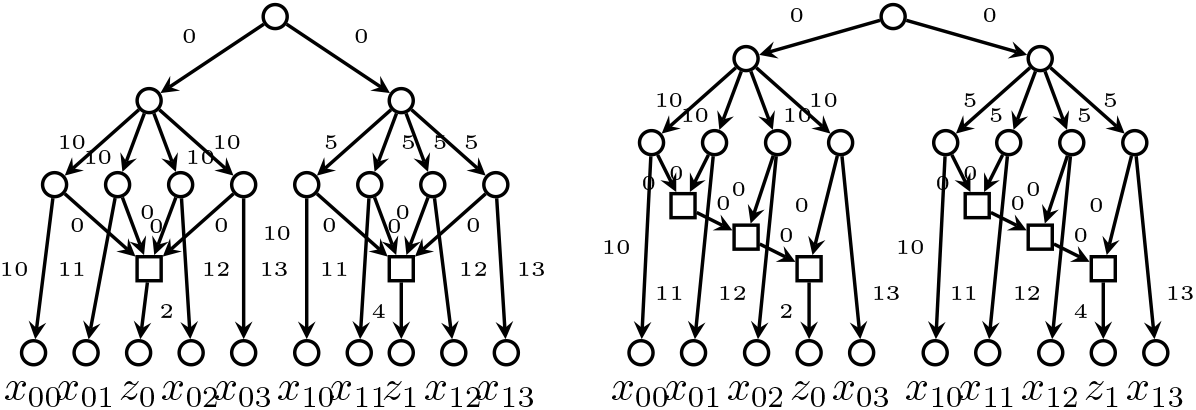
Networks from Example 21.

In both cases we have the following optimal sets of leaves:

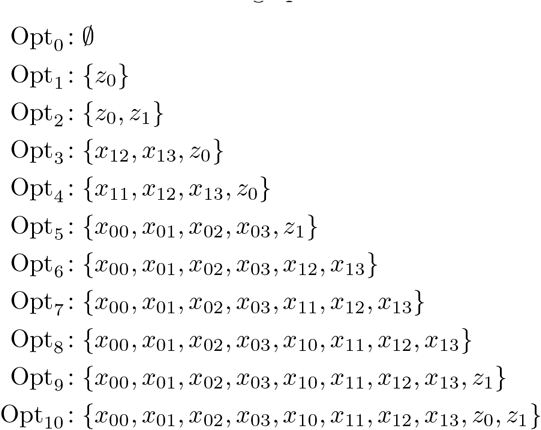

In the case of the semi-4-ary level-1 network we have

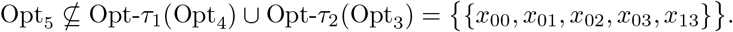

However, for the level-3 network, Opt_*m*_ = Opt-*τ*_1_(Opt_*m*−1_) and 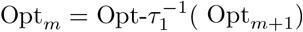 for all 1 ⩽ *m* ⩽ *n* = 10 and 0 ⩽ *m < n* respectively. This should not be surprising, because in Example 6 we showed that for this network (although with different weights) we can always find an rPSD-improving pair (*A, B*) with |*A*| − |*B*| = 1, hence the first case in the proof of Proposition 19 could always be chosen and prove that Opt_*m*_ ⊆ Opt-*τ*_1_(Opt_*m*−1_).

As we mention in Example 21 below, if some network has some Opt_*m*_ not included into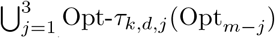, then that network has two sets of leaves *X,X*_*′*_with *m* = |*X*| = |*X*_*′*_| + 1 and no rPSD-improving pairs (*A, B*) with |*A*| − |*B* | *<* 3. Otherwise, if we could always find some rPSD-improving pair with |*A*| − |*B* | *<* 3, the proof of Propositions 19 and 20 would never need to explore the cases *j*_1_ ∈ {3, 4} and thus obtain a similar result to Proposition 17. In Example 22 we show a semi-5-ary network that has *X* = {*x*_00_, …, *x*_04_, *z*_1_} and *X*^*′*^ = {*x*_10_, …, *x*_13_, *z*_0_}with only rPSD-improving pairs with |*A| − |B|* ⩾ 3, yet the obvious greedy algorithm would still work in this network. In contrast, we have not found any semibinary level-3 network that has some *X, X*^*′*^ ⊆ Σ with |*X*| = |*X*^*′*^| + 1 and no rPSD-improving pair (*A, B*) with |*A*| − |*B*| = 1.

### Example 22.

The semi-5-ary level-1 network in Figure 7, analogous to the semi-4-ary network from Example 21, similarly has 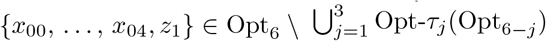 but still, for all 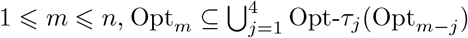.

**Figure 7:**
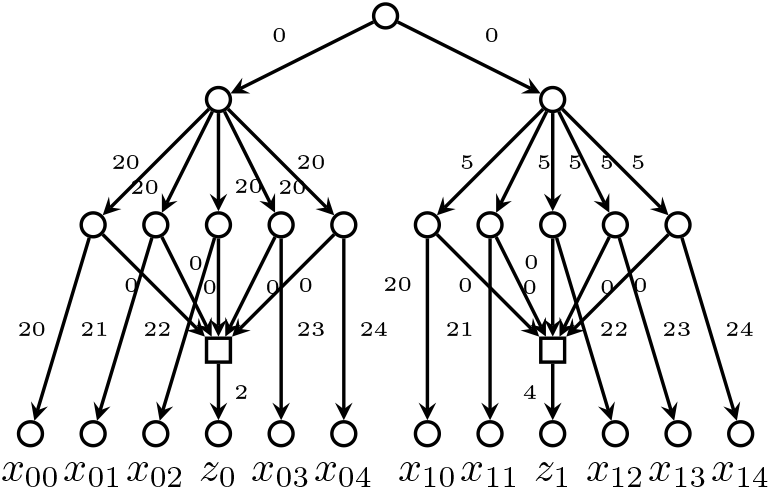
Network from Example 22.

## 6 Conclusions

PD on phylogenetic trees satisfies the strong exchange property that guarantees that the sum of the PD values of two sets of leaves of different cardinalities can always be increased by moving an element from the largest set to the smallest one, but rPSD does not longer satisfy this exchange property even for galled trees. In this paper we have generalized this exchange property to rPSD on phylogenetic networks of bounded level an reticulations’ in-degree, showing that a similar results holds if we allow more involved interchanges of leaves’ subsets. Our final goal was then to use this generalized exchange property to find a polynomial time greedy algorithm for the optimization of rPSD on phylogenetic networks of bounded level and in-degree of reticulations. We have ultimately failed in this goal. We have indeed shown that the generalized exchange property entails such a greedy algorithm for semibinary level-2 networks and semi-3-ary level-1 networks (and sheds new light on the structure of the families of rPSD-optimal sets Opt_*m*_ on galled trees) but it cannot be used, as it stands, to obtain such an algorithm on more complex networks. However, we have not been able to find examples of semibinary level-3 networks or semi-4-ary level-1 networks where the greedy algorithm fails: it is simply that the generalized exchange property alone is not enough to prove its correctness.

Finally, it is important to point out that just like the rPSD optimization problem itself, testing counterexamples is computationally expensive too. While the greedy algorithm runs in polynomial time, finding whether Opt_*m*_ can be obtained from some Opt_*m − j*_ still requires calculating Opt_*m*_ by brute force, and testing whether the exchange property holds for a certain subset of 𝒮_*k,d*_ where |*A*| − |*B*| *< j* also requires testing all subsets *X, X*^*′*^ ⊆Σ. All of these are exponential operations, hence trying even slightly larger examples can dramatically increase the runtime of the test.

## Acknowledgements

This research was partially supported by the Spanish Ministry of Economy and Competitiveness and European Regional Development Fund project PGC2018-096956-B-C43 (MINECO/FEDER) and the grant PID2021-126114NB-C44 funded by MCIN/AEI/10.13039/501100011033 and by “ERDF A way of making Europe.”

## Appendix: Proof of Corollary 9

We first prove a refinement of Lemma 5.

### Lemma 23.

*Let* ℬ *be a semi-d-ary level-1 blob with exit reticulation H and X, X*^*′*^ *two multisets of nodes of* ℬ *satisfying the following properties:*

i. |*X*^*′*^| *<* |*X*|.
ii. *For each v* ∈ *V* (ℬ), *if m*_*X*′_ (*v*) *< m*_*X*_ (*v*), *then m*_*X*_ (*v*) = 1.
iii. *H* ∈ *X* ∪ *X*^*′*^.

*Then there exist two sets A, B of nodes of* ℬ *such that:*

1. |*B*| = 0, |*A*| = 1 *or* |*B*| = 1 < |*A*| ⩽ *d*.
2. *A* ⊆ Supp *X* \ Supp *X*′.
3. *B* ⊆ {*H*} ∩ (Supp *X*′ \ Supp *X*).
4. ↑*X* ∩ ↑*X*′ ⊆ ↑*τ*_*A,B*_(*X*) ∩ ↑*τ*_*B,A*_(*X*′).

*Proof*. Using the same notations as in Lemma 5, observe that the equation (2) holds identically in this case, i.e.,

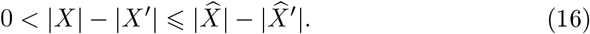

In addition, ℰ= {*H*}. Now consider the following cases:

a. If there exists some 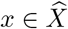 with a strict descendant in *X*, then *A* = {*x*}, *B* = ∅ satisfy the required conditions as proved in case **(a)** of Lemma 5.
b. If *H* ∈ *X* and no 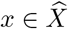 has any strict descendant in *X*, then 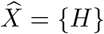, and then 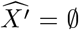 (by Eqn. (16) and ↑*X*^*′*^ = ↑(*X* | {*H*}) because, since 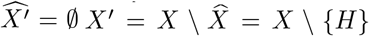,. Then, taking *A* = {*H*} and *B* = ∅ we have that properties (1) to (4) trivially hold.
c. If *H* ∉ *X* and no 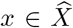 has any strict descendant in *X*, then, on the one hand, *H* belongs to Supp *X*^*′*^*∖* Supp *X* by condition *(iii)*, and hence 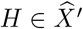, and, on the other hand, 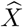 is an independent set of at most *d* nodes (because there are at most *d* different paths from the root to *H*).

For brevity, let *X*^*′′*^ denote the full sub-multiset of *X*^*′*^ supported on Supp *X*^*′*^ ∖ {*H*}. By Eqn. (16),

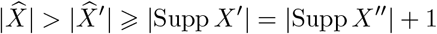

and thus 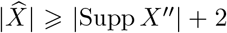. Now, since ℬ does not contain internal reticulations and the nodes in 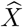 are independent, each node in *X*^*′′*^ has at most one ancestor in 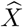. This implies that 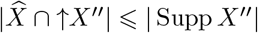 and hence

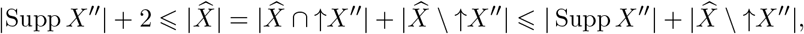

which implies 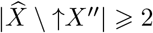.

Take then 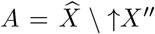 and *B* = {*H*}. As we have just seen, 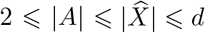. Thus, *A, B* satisfy properties (1)–(3). As far as (4) goes, that is,

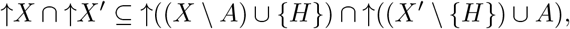

observe that, since *H* is the only exit reticulation of ℬ, ↑*H* = *V* (ℬ) and hence ↑*X*^*′*^ = ↑((*X \ A*) ∪ {*H*}) = *V* (*ℬ*). Therefore, we actually must prove that

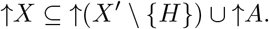

We must consider two cases.

- If *m*_*X′*_ (*H*) *>* 1, then *H ∈ X*^*′*^*∖* {*H*} and thus ↑ (*X*^*′*^*∖* {*H*}) = *V* (ℬ) and the desired inclusion is obvious.
- If *m*_*X′*_ (*H*) = 1, then *X*^*′*^ *∖* {*H*} = *X*^*′′*^, and in this case

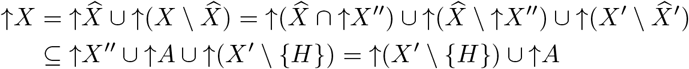

as desired.

This finishes the proof of case **(c)**.

We can proceed now with the proof of Corollary 9. Arguing as in the proof of Theorem 6, we can assume that *N* is at-most-bifurcating and in particular that no node in *N* is the split node of more than one blob. Notice moreover that now each blob has only one reticulation: its exit reticulation.

We follow the proof of Theorem 6 by induction on the number *α* of edges of the network. The base case *α* = 0 is again obvious, and thus we must only consider the inductive step. So, let *N* be a semi-*d*-ary level-1 weighted phylogenetic network on Σ with more than one edge, and let *X, X*^*′*^*⊆* Σ with |*X*|^*′*^ *< X*. If *X* = 1 the exchange property is trivially satisfied taking (*A, B*) = (*X*,) 𝒮 _0_(Σ), so we assume | *X* | ⩾ 2. As in the proof of the aforementioned theorem, we shall denote its clusters *C*_*N*_ (*v*) by simple *C*(*v*).

Arguing as in cases **(a), (b)** and **(c)** in the proof of Theorem 6, we can assume that:

i. *The root r is the split node of a single, semi-d-ary blob ℬ*.
ii. *For every node v in ℬ, if v has a child* 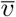 *outside ℬ such that* 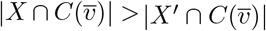, *then* 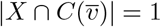 *and* 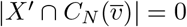.
iii. *X* ∩ *C*(*H*) *≠* ∅ or *X*^*′*^ ∩ *C*(*H*) *≠* ∅.

We use henceforth the same notations as in point **(d)** in the proof of that theorem. The hypotheses of Lemma 23 are satisfied by 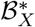 and 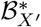. Then, there exist two sets *ℬ* _*A*_, *ℬ* _*B*_ ⊆ *V* (*ℬ*) such that:

1. | *ℬ* _*B*_| = 0 and | *ℬ* _*A*_| = 1, or | *ℬ* _*B*_| = 1 *<* | *ℬ* _*A*_| ⩽ *d*.
2. *ℬ* _*A*_ ⊆ Supp 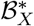 ∖ Supp 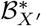.
3. *B*_*B*_ ⊆ {*H*} ∩ (Supp 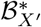 *∖* Supp 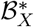).
4. 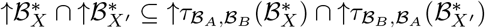.

Now take 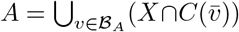 and, if *ℬ*_*B*_ = {*H*}, choose any *b* ∈ *X*^*′*^∩*C*(*H*) and take *B* = {*b*}, while if *ℬ*_*B*_ = ∅, take *B* = ∅. Notice that, by (1), if *B* = ∅ then |*B*_*A*_| = 1 and hence |*A*| = 1, while, if |*B*| = 1, then *H* ∈ Supp 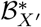 *∖*Supp 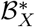 and hence *X* ∩ *C*(*H*) = ∅.

These sets satisfy |*A*| = |ℬ_*A*_|, |*B*| = |ℬ_*B*_| and thus (*A, B*) ∈ 𝒮_*d*_(Σ). Now, arguing as in Theorem 6, we have 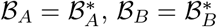and

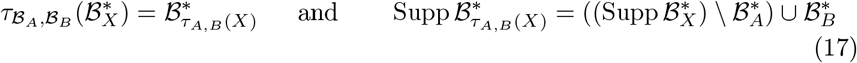

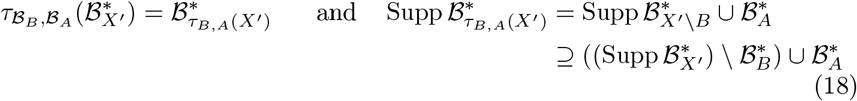

(where the inclusion in (18) is not an equality if *B*_*B*_ = {*H*} and 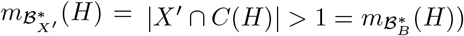 and, still arguing as in Theorem 6,

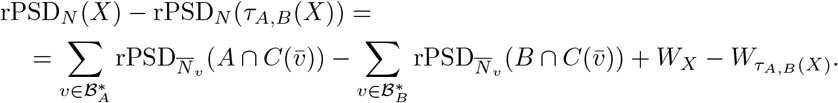

Now, the difference between (18) here and (8) in the proof of Theorem 6 makes dealing with rPSD_*N*_ (*τ*_*B,A*_(*X*^*′*^)) − rPSD_*N*_ (*X*^*′*^) different here from in the aforementioned proof. In the current situation, we have that

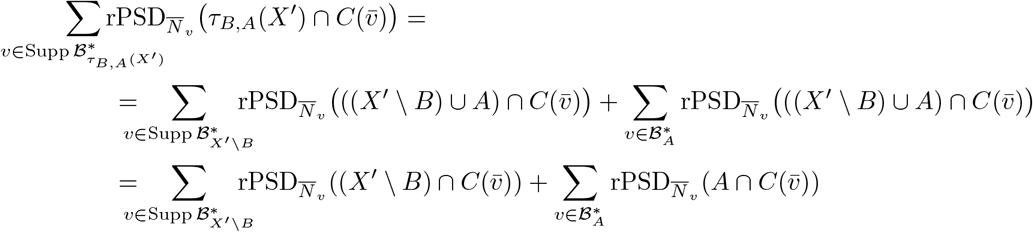

(because, if 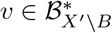, then 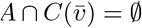, and if 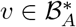, then 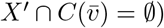;

and

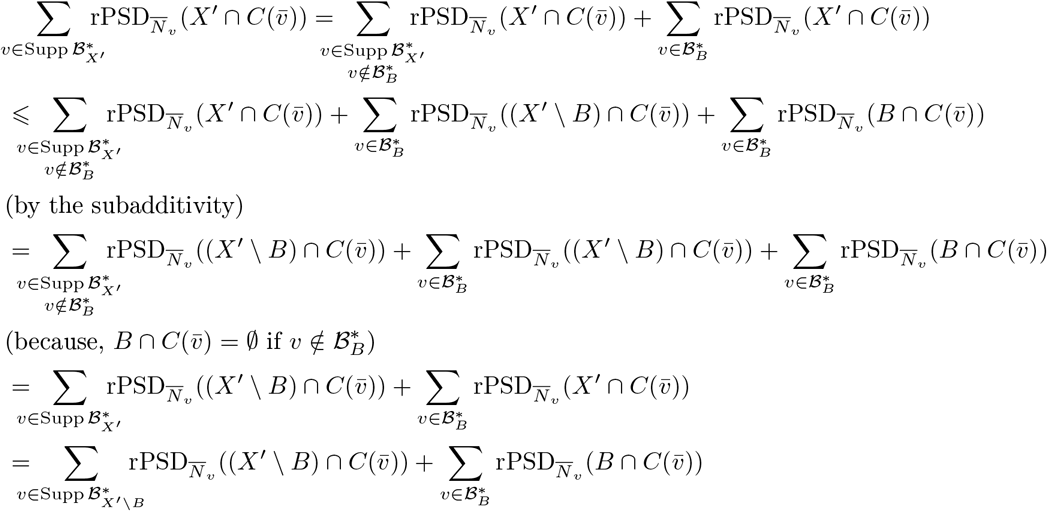

(because 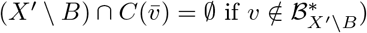. Therefore

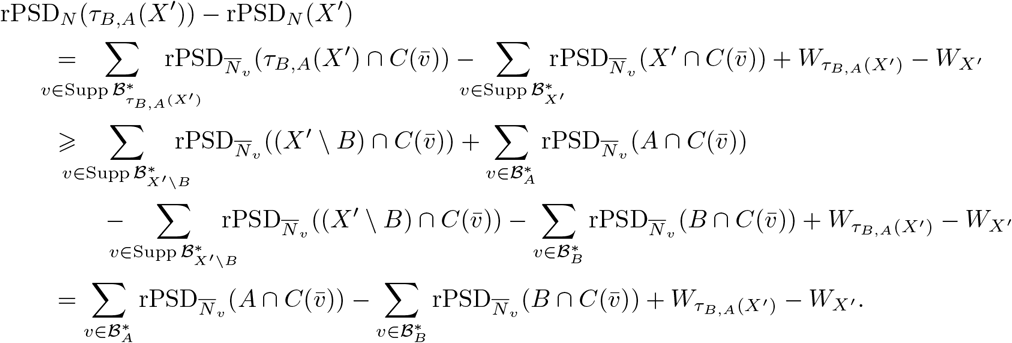

Then, as in Theorem 6, we conclude that

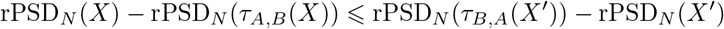

if

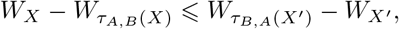

and this last inequality is deduced from property (4) as in the proof of Theorem 6.

## References

[1] M. Baroni, C. Semple, M. Steel. “Hybrids in real time.” Systematic Biology 55 (2014) 46–56.

[2] R. G. Beiko. “Telling the whole story in a 10,000-genome world.” Biology direct, 9 (2014), 18.

[3] M. Bordewich, C. Semple, A. Spillner. “Optimizing phylogenetic diversity across two trees.” Applied Mathematics Letters, 22 (2009), 638–641.

[4] M. Bordewich, C. Semple, K. Wicke. “On the Complexity of Optimising Variants of Phylogenetic Diversity on Phylogenetic Networks.” Theoretical Computer Science 917 (2022), 66–80.

[5] G. Cardona, A. Mir, L. Rotger, F. Rosselló, D. Sánchez, “Cophenetic metrics for phylogenetic trees, after Sokal and Rohlf.” BMC Bioinformatics (2013) 14:3

[6] G. Cardona, F. Rosselló, G. Valiente. “Comparison of tree-child phylogenetic networks.” IEEE/ACM Transactions on Computational Biology and Bioinformatics (TCBB) 6 (2009), 552–569.

[7] O. Chernomor, S. Klaere, et al. “Split diversity: measuring and optimizing biodiversity using phylogenetic split networks.” In [27] (2016), 173–195.

[8] T. M. Coronado, G. Riera, F. Rosselló. “The Fair Proportion is a Shapley Value on phylogenetic networks too.” In: Enjoying Natural Computing (C. Graciani, A., Riscos-Núñez et al eds.) Springer (2018), 77–87.

[9] W. F. Doolittle. “Phylogenetic classification and the universal tree.” Science 284 (1999), 2124–2128.

[10] D. Faith. “Conservation evaluation and phylogenetic diversity.” Biological Conservation 61 (1992), 1–10.

[11] M. Fuchs, E. Y. Jin. “Equality of Shapley value and fair proportion index in phylogenetic trees.” Journal of Mathematical Biology 71 (2015), 1133–1147.

[12] P. Gambette, V. Berry, C. Paul. The structure of level-k phylogenetic networks. In: Annual Symposium on Combinatorial Pattern Matching. Springer (2009), 289–300.

[13] K. J. Gaston. “Species richness: measures and measurements”. In: Biodiversity: a biology of numbers and differences (K. J. Gaston ed.), Blackwell Science (1996) pp. 77–113.

[14] R. Gumbs, C. L. Gray et al. “The EDGE2 protocol: Advancing the prioritisation of Evolutionarily Distinct and Globally Endangered species for practical conservation action.” PLoS Biology 21 (2023), e3001991.

[15] D. Gusfield, S. Eddhu, C. Langley. “Optimal, efficient reconstruction of phylogenetic networks with constrained recombination.” Journal of Bioinformatics and Computational Biology 2 (2004) 173–213.

[16] D. Gusfield, V. Bansal, et al. “A decomposition theory for phylogenetic networks and incompatible characters.” Journal of Computational Biology 14 (2007), 1247–1272.

[17] C.-J. Haake, A. Kashiwada, F. E. Su. “The Shapley value of phylogenetic trees.” Journal of Mathematical Biology 56 (2008), 479–497.

[18] D. Huson, D. Bryant. “Application of phylogenetic networks in evolutionary studies.” Molecular biology and evolution, 23 (2006), 254–267.

[19] D. Huson, R. Rupp, C. Scornavacca. Phylogenetic Networks: Concepts, Algorithms and Applications. Cambridge University Press (2010).

[20] N. Isaac. S. T. Turvey et al. “Mammals on the EDGE: Conservation priorities based on threat and phylogeny.” PLoS ONE 2 (2007), e296.

[21] J. Jansson, W.-K. Sung. “Inferring a level-1 phylogenetic network from a dense set of rooted triplets.” Theoretical Computer Science 363 (2006) 60–68.

[22] L. Jetten, L. van Iersel. “Nonbinary tree-based phylogenetic networks.” IEEE/ACM Transactions on Computational Biology and Bioinformatics 15 (2018), 205–217.

[23] E. Kolbert. The Sixth Extinction. An Unnatural History. Henry Holt and Company (2014).

[24] J. A. McNeely, K. R. Miller, et al. Conserving the world’s biological diversity. International Union for conservation of nature and natural resources (1990).

[25] G. Nemhauser, L. Wolsey, M. Fisher. “An analysis of approximations for maximizing submodular set functions–I.” Mathematical Programming 14 (1978), 265–294.

[26] F. Pardi, N. Goldman. “Species choice for comparative genomics: Being greedy works.” PLoS Genetics 1 (2005), e71.

[27] R. Pellens, P. Grandcolas eds. Biodiversity conservation and phylogenetic systematics: preserving our evolutionary heritage in an extinction crisis Springer Nature (2016).

[28] H. P. Possingham, S. Andelman et al. “Limits to the use of threatened species lists.” Trends in Ecology & Evolution 17 (2002), 503–507

[29] D. Redding, A. Mooers. “Incorporating evolutionary measures into conservation prioritization.” Conservation Biology 20 (2006), 1670–1678.

[30] G. Riera. Theoretical Models and Computational Techniques for the Analysis of Microbial Communities. PhD Thesis, UIB (2023).

[31] C. Solís-Lemus, C. Ané. Inferring phylogenetic networks from gene order data. Molecular biology and evolution, 33 (2016), 1207–1219.

[32] A. Spillner, B. T. Nguyen, V. Moulton. “Computing phylogenetic diversity for split systems.” IEEE/ACM Transactions on Computational Biology and Bioinformatics 5 (2008), 235–244.

[33] M. Steel. “Phylogenetic diversity and the greedy algorithm.” Systematic Biology 54 (2005), 527–529.

[34] M. Steel. Phylogeny: Discrete and random processes in evolution. SIAM (2016).

[35] C. M. Tucker, M. W. Cadotte et al. “A guide to phylogenetic metrics for conservation, community ecology and macroecology.” Biological Reviews, 92 (2017), 698–715.

[36] M. Vellend, W. Cornwell, et al. “Measuring phylogenetic biodiversity”. In: Biological diversity: Frontiers in measurement and assessment (A. E. Magurran and B. J. McGill, eds), Oxford University Press (2010), 194–207.

[37] L. Volkmann, I. Martyn et al. “Prioritizing populations for conservation using phylogenetic networks.” PloS one 9 (2014), e88945.

[38] K. Wicke, M. Fischer. “Phylogenetic diversity and biodiversity indices on phylogenetic networks.”. Mathematical Biosciences 298 (2018), 80–90.

[39] Y. Yu, J. Dong, K. J. Liu. “Bayesian estimation of species networks from multilocus data.” Molecular biology and evolution, 31 (2014), 1032–1043.

[40] A. Zhukova, L. Blassel et al. “Origin, evolution and global spread of SARS-CoV-2.” Comptes Rendus. Biologies 344 (2021), 57–75

